# Genetic dissection reveals the role of Ash1 domains in counteracting Polycomb repression

**DOI:** 10.1101/675173

**Authors:** Eshagh Dorafshan, Tatyana G. Kahn, Alexander Glotov, Mikhail Savitsky, Yuri B. Schwartz

## Abstract

Antagonistic functions of Polycomb and Trithorax proteins are essential for proper development of all metazoans. While the Polycomb proteins maintain the repressed state of key developmental genes, the Trithorax proteins ensure that these genes stay active in cells where they have to be expressed. Ash1 is the Trithorax protein that was proposed to counteract Polycomb repression by methylating lysine 36 of histone H3. However, recently it was shown that genetic replacement of *Drosophila* histone H3 with the variant that carried Arginine instead of Lysine at position 36 did not impair the ability of Ash1 to counteract Polycomb repression. This argues that Ash1 counteracts Polycomb repression by methylating, yet unknown, non-histone proteins. To find these substrates, one may need to look beyond the function of the Ash1 histone methyltransferase SET domain at other evolutionary conserved parts of the protein that received little attention. Here we used *Drosophila* genetics to demonstrate that Ash1 requires each of the BAH, PHD and SET domains to counteract Polycomb repression, while AT hooks are dispensable. Our findings argue that, *in vivo*, Ash1 acts as a multimer. Thereby, it can combine the input of the SET domain and PHD-BAH cassette residing in different peptides. Finally, using new loss of function alleles, we show that zygotic Ash1 is required to prevent erroneous repression of homeotic genes.

## Introduction

Embryonic development is controlled by genes encoding morphogenes and transcriptional regulators. These genes need to be switched on in correct cells at appropriate time and their expression, or lack thereof, has to be maintained as the embryo grows and cells continue to divide. Polycomb and Trithorax systems are critical to maintain the expression status of developmental genes (Schwartz and Pirrotta 2007; Schuettengruber *et al*. 2017). Protein components of the Polycomb system assemble in complexes that act as epigenetic repressors. While mechanistic details of the repression are still being worked out, studies in fruit flies indicate that tri-methylation of Lysine 27 of histone H3 (H3K27) by one of the Polycomb complexes (PRC2) is essential for the process (Pengelly *et al*. 2013; McKay *et al*. 2015; Coleman and Struhl 2017; Laprell *et al*. 2017).

The Trithorax system counteracts Polycomb repression to ensure that developmental genes repressed by Polycomb complexes in certain cell types are not erroneously shut down in cells where they have to remain active. Genetic evidence from studies on *Drosophila melanogaster* indicate that the Trithorax (Trx) and Absent, small, or homeotic discs 1 (Ash1) proteins are critical components of the system (Poux *et al*. 2002; Klymenko and Muller 2004). Of the two proteins, Ash1 is particularly interesting. Unlike Trx, which binds Polycomb regulated genes regardless of their transcriptional state (Schuettengruber *et al*. 2009; Schwartz *et al*. 2010), Ash1 binds and forms extensive chromatin domains exclusively when Polycomb regulated genes are transcriptionally active and Polycomb repression is impaired (Schwartz *et al*. 2010; Kharchenko *et al*. 2011; Huang *et al*. 2017). Trx and Ash1 are incorporated in distinct complexes whose composition is yet to be fully characterized (Petruk *et al*. 2001; Mohan *et al*. 2011; Huang *et al*. 2017; Schmahling *et al*. 2018) but both have SET domains and can methylate histone H3 (Smith *et al*. 2004; Tanaka *et al*. 2007; An *et al*. 2011; Yuan *et al*. 2011; Tie *et al*. 2014). *In vitro* experiments indicate that the histone methyltransferase activity of PRC2 is inhibited by prior methylation of histone H3 tail at Lysine 4 (H3K4) or Lysine 36 (H3K36) (Schmitges *et al*. 2011; Yuan *et al*. 2011; Voigt *et al*. 2012), the sites that can be methylated by Trx and Ash1. From this, it was proposed that Trx and Ash1 counteract Polycomb repression by inhibiting PRC2 catalytic activity via H3K4 and H3K36 methylation (Schmitges *et al*. 2011; Yuan *et al*. 2011; Voigt *et al*. 2012). Naturally, most recent studies of Trx and Ash1 have focused on mechanistic details of histone H3 methylation by their SET domains (Dorighi and Tamkun 2013; Tie *et al*. 2014; Huang *et al*. 2017; Schmahling *et al*. 2018; Dorafshan *et al*. 2019; Hou *et al*. 2019; Lee *et al*. 2019).

Although the “PRC2-inhibition” model nicely explains the antagonistic relation between Polycomb and Trithorax systems, it is at odds with results of *in vivo* studies. Thus, experiments of Hödl and Basler (Hodl and Basler 2012), as well as our recent work (Dorafshan *et al*. 2019), showed that genetic replacement of *Drosophila* histone H3 with variants that carry Arginine instead of Lysine at positions 4 or 36 does not impair the ability of Trx and Ash1 to counteract Polycomb repression. Since intact SET domains of both proteins are required to antagonize the repression (Stassen *et al*. 1995; Dorafshan *et al*. 2019), this argues that Trx and Ash1 counteract Polycomb system by methylating, yet unknown, non-histone proteins. To find these substrates, it may be time to look more carefully at other evolutionary conserved domains of these proteins which, so far, received little attention. Are these domains critical for Trx and Ash1 function? Do they act in concert or as separate modules? Which of them, if any, are critical to bind chromatin? Here we used the power of *Drosophila* genetics to address these questions for the protein domains of Ash1. Our experiments demonstrate that Ash1 requires each of the BAH, PHD and SET domains to counteract Polycomb repression, while AT hooks are dispensable. Our findings argue that, *in vivo*, Ash1 acts as a multimer and can combine the input of SET and PHD or BAH domains residing in different monomers. Finally, using new loss of function *ash1* alleles, we demonstrate that its zygotic function is required to prevent erroneous repression of homeotic genes.

## Materials and methods

### Plasmid construction

Ash1 expressing constructs were assembled by Gateway LR recombination (Invitrogen) between an entry construct containing corresponding OneSTrEP-tagged CDS and the destination vector pWattB-Ubi-DEST. To generate entry constructs with truncated Ash1 CDS, corresponding deletions were introduced into the entry construct containing full-length Ash1 CDS (pENTR1A-OneSTrEP-Ash1FL) as described below. The pENTR1A-OneSTrEP-Ash1FL and pWattB-Ubi-DEST constructs are described in (Dorafshan *et al*. 2019).

To generate pENTR1A-OneSTrEP-Ash1ΔSET construct, deltaSET_AB and deltaSET_CD fragments, flanking the SET domain, were amplified from pENTR1A-OneSTrEP-Ash1FL using primers deltaSET_A, deltaSET_B, and deltaSET_C, deltaSET_D. pENTR1A-OneSTrEP-Ash1FL was digested with BstZ17I and SphI and deltaSET_AB and deltaSET_CD fragments were introduced to the linear vector using InFusion (Clontech). pENTR1A-OneSTrEP-Ash1ΔSET plasmid was sequenced using ASH1_seq7 and ASH1_seq8 primers.

To construct pENTR1A-OneSTrEP-Ash1ΔPHD, the upstream deltaPHD_AB and downstream deltaPHD_CD fragments, flanking the PHD domain, were amplified from pENTR1A-OneSTrEP-Ash1FL using primers deltaPHD_A, deltaPHD_B and deltaPHD_C, deltaPHD_D. After this, pENTR1A-OneSTrEP-Ash1FL was digested with KpnI and SphI and deltaPHD_AB and deltaPHD_CD fragments were introduced to the linear vector using InFusion. pENTR1A-OneSTrEP-Ash1ΔPHD plasmid was sequenced using ASH1_seq8, ASH1_seq9, and ASH1_seq12 primers.

To generate pENTR1A-OneSTrEP-Ash1ΔBAH construct, two fragments (upstream deltaBAH_AB and downstream deltaBAH_CD) flanking BAH domain were amplified from pENTR1A-OneSTrEP-Ash1FL using primers deltaBAH_A, deltaBAH_B and deltaBAH_C, deltaBAH_D. pENTR1A-OneSTrEP-Ash1FL was digested with ClaI and XhoI restriction enzymes. Linear vector (8.7 kb) was extracted from the gel and used in the InFusion reaction together with fragments deltaBAH_AB and deltaBAH_CD. The resulted pENTR1A-OneSTrEP-Ash1ΔBAH construct was sequenced using ASH1_seq13 and ASH1CN_Cfwd primers.

To obtain pENTR1A-OneSTrEP-Ash1ΔAT construct, all three AT hooks were deleted sequentially. Two fragments (deltaAT_EF upstream and deltaAT_GH downstream) flanking the second AT hook were amplified from pENTR1A-OneSTrEP-Ash1FL using primers deltaAT_E, deltaAT_F and deltaAT_G, deltaAT_H. pENTR1A-OneSTrEP-Ash1FL was digested with SacII and BstZ17I restriction enzymes. Linear vector (9 kb) was extracted from the gel and used in the InFusion reaction together with fragments deltaAT_EF and deltaAT_GH to yield the pENTR1A-OneSTrEP-Ash1ΔAT2 construct. Next, two fragments (deltaAT_EI upstream and deltaAT_HJ downstream) flanking the third AT hook were amplified from pENTR1A-OneSTrEP-Ash1ΔAT2 plasmid using primers deltaAT_E, deltaAT_I and deltaAT_H, deltaAT_J. pENTR1A-OneSTrEP-Ash1ΔAT2 was digested with SacII and BstZ17I restriction enzymes. Linear vector (9 kb) was extracted from the gel and used in the InFusion reaction together with fragments deltaAT_EI and deltaAT_HJ to obtain pENTR1A-OneSTrEP-Ash1ΔAT23. Finally, two fragments (deltaAT_AB upstream and deltaAT_CD downstream) flanking the first AT hook were amplified from pENTR1A-OneSTrEP-Ash1ΔAT23 using primers deltaAT_A, deltaAT_B and deltaAT_C and deltaAT_D. pENTR1A-OneSTrEP-Ash1ΔAT23 was digested with SacI. Linear vector (9 kb) was extracted from the gel and used in the InFusion reaction together with fragments deltaAT_AB and deltaAT_CD to get the final pENTR1A-OneSTrEP-Ash1ΔAT construct. The pENTR1A-OneSTrEP-Ash1ΔAT2 and pENTR1A-OneSTrEP-Ash1ΔAT23 plasmids were sequenced using ASH1_seq4, ASH1_seq5, ASH1_seq6, and ASH1_seq7 primers. The pENTR1A-OneSTrEP-Ash1ΔAT was further sequenced using ASH1_seq3 and ASH1_seq14 primers.

To generate pWattB-U6.2-2targets plasmid, the target sequences for CRISPR/Cas9-mediated cleavage (SET-CRSP-1 and poSET-CRSP-1) were selected using Cas9 Target Finder software (http://www.shigen.nig.ac.jp/fly/nigfly/cas9/index.jsp). SET-CRSP-1 is located within *ash1* SET domain and poSET-CRSP-1 is located after the *ash1* postSET domain 243bp apart. Corresponding pairs of complementary oligonucleotides (SET-CRSP-1.1 and SET-CRSP-1.2) and (poSET-CRSP-1.1 and poSET-CRSP-1.2) were ordered from Sigma-Aldrich and annealed to obtain double stranded SET-CRSP-1 and poSET-CRSP-1 fragments. The DNA of the pBFv-U6.2B and pBFv-U6.2 plasmids (kind gifts from Dr. Maria Kim) was digested with BbsI restriction enzyme (Fermentas, Cat # ER1011) and the linear products ligated with SET-CRSP-1 and poSET-CRSP-1 fragments resulting in pBFv-U6.2B-SET and pBFv-U6.2-poSET constructs, respectively. pBFv-U6.2-poSET construct was digested with EcoRI and NotI and the resulting 517bp fragment (U6.2-poSET) isolated and ligated with the pBFv-U6.2B-SET construct, digested with the same enzymes. This yielded the pBFv-U6.2-2targets constructs. Following this, the DNA of the pBFv-U6-2B_2targets construct was digested with SacI. The resulting 1.2kb SacI fragment (containing U6-2B_2targets) was blunt-end ligated with DNA of the pWattB plasmid (Savitsky *et al*. 2016) digested with XhoI, which yielded pWattB-U6.2-2targets construct. For corresponding oligonucleotide sequences see Table S1.

### Fly strains

*w*; ash1^22^,P{w^+mW.hs^=FRT(w^hs^)}2A/TM6C,Sb^1^,Tb^1^* (Bloomington Drosophila Stock Center, 24161), *w^1118^; ash1^Df(3L)Exel9011^/TM6B,Tb^1^* flies (Bloomington Drosophila Stock Center, 7945), and *w^1^; ash1^21^/TM3,Ser* (Steffen *et al*. 2013) were re-balanced over *TM3,Ser,e,Act-GFP^+mW^* balancer*. Oregon R* flies (kind gift from Dr. Jan Larsson) were used as wild-type in all experiments unless stated otherwise. *y^2^,cho^2^,v^1^; attP40{nos-Cas9}/CyO* fly strain (Kondo and Ueda 2013) was used as a source of Cas9. *w^1^; If/CyO; MKRS/TM6, Tb* was used to balance CRISPR/Cas9-edited chromosomes.

### Fly transformation

All constructs were injected in pre-blastoderm embryos by BestGene Inc. The Ash1 expressing (full-length and truncated) transgenes were injected in the *y^1^ M{vas-int.Dm}ZH-2A w*; M{3xP3-RFP.attP}ZH-51C* (24482) strain. The pWattB-U6.2-2targets construct was injected in the *y^1^,w^67c23^; P{CaryP}attP40* strain.

### CRIPSR/Cas9-mediated genome editing

The editing strategy described in (Kondo and Ueda 2013) was used to generate deletions within Ash1 SET domain. Flies of the *y^1^ w^67c23^; pWattB-U6.2-2targets* strain, expressing two gRNA, were crossed to *y^2^,cho^2^,v^1^; attP40{nos-Cas9}/CyO*. Resulting *attP40{nos-Cas9}/pWattB-U6.2-2targets; +/+* females were crossed to *w^1^; If/CyO; MKRS/TM6,Tb* and the progeny individually screened for editing events by PCR with ash1_seq8 and deltaSET_D primers which amplify 499bp fragment from unedited chromosomes and 262bp fragment from chromosomes with precise deletion.

### Polytene chromosome preparation and immunostaining

Salivary glands were dissected from 3rd instar larvae, preparation and immunostaining of polytene chromosomes was performed as described elsewhere (Eggert *et al*. 2004). Images were taken with Zeiss Apotome Microscope equipped with Plan-Apochromat 63×/1.40 oil DIC M27 objective, filters set (63HE for red channel, 38HE for green channel and 49 for DAPI) and AxioCam MR R3 camera. Images were processed with ZenPro software (Zeiss) and mounted in ImageJ. For the list of antibodies see Table S2.

### Fly cuticle preparation and embryo immunostaining

Flies were boiled in 10% KOH for 10 minutes, incubated in distilled water for 30 minutes, and dehydrated in 70% ethanol and 99% ethanol for 10 minutes each. Ethanol was removed and replaced with glycerol. Cuticles were incubated for 30 minutes in glycerol, then dissected under the stereo microscope and mounted on glass slide in glycerol. Embryos were immunostained as described in (Dorafshan *et al*. 2019). For the list of antibodies see Table S2.

### Chromatin immunoprecipitation (ChIP)

ChIP and qPCR analysis were performed as described (Kahn *et al*. 2016) except that crosslinked material was sonicated in 4ml of 10mM Tris-HCl pH8.0, 1mM EDTA pH8.0 for 45 minutes with Branson 450 Digital Sonifier (45 cycles of 20 sec ON – 40 sec OFF). The isolated ChIP material was re-suspended in 400 µl of DNase free water and 4 µl were used for each quantitative PCR reaction. The antibodies used are listed in Table S2 and the ChIP amplicons are listed in Table S3.

### Western blot

Nuclear extracts from whole 3rd instar larvae were separated on a 6% SDS-PAGE and blotted to PVDF membrane for 3 hours at 200mA. The same extracts were separated on a 15% SDS-PAGE and stained with Coomassie to be used as loading control. Primary and secondary antibodies were diluted in 1xPBS with 1% BSA and 0,05% Tween-20. For the list of antibodies, see Table S2.

## Results

To investigate how individual domains enable Ash1 to counteract Polycomb repression, we made an array of transgenic constructs (Figure 1). Each construct contained either the full length Ash1 Open Reading Frame (ORF) or truncated Ash1 ORFs lacking specific domains and *attB* site for phiC31-mediated site-specific recombination (Bischof *et al*. 2007). All constructs were driven by *Ubiquitin* (*Ubi-p63E*) promoter (Butcher *et al*. 2004) and integrated in the same *51C* genomic site on chromosome 2L. The latter helped to reduce the variability in the expression of transgenic constructs providing identical chromatin environment.

**Figure 1.**
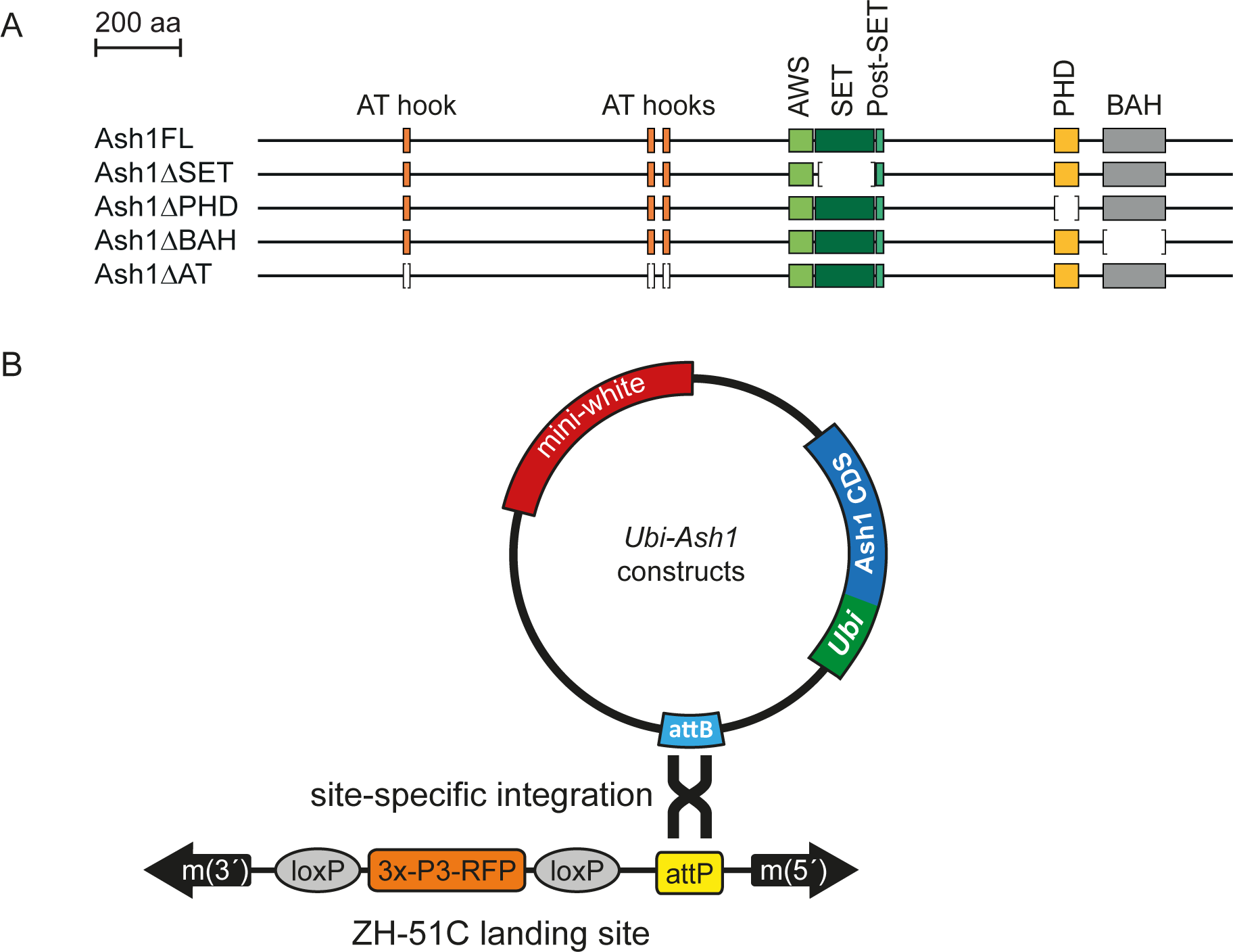
Generation of *Ash1* transgenes. **(A)** Transgenic Ash1 variants. Four different mutant variants were generated starting from the full-length *Ash1* cDNA (Ash1FL). SET (dark green box), PHD (yellow box) and BAH (grey box) domains were deleted in Ash1ΔSET, Ash1ΔPHD and Ash1ΔBAH variants, respectively. In the Ash1ΔAT variant all three AT hooks (orange boxes) were deleted. Brackets show corresponding deleted regions. **(B)** All *Ash1* Open Reading Frames from above were cloned in a vector containing *Ubi-p63E* promoter, an attB integration site, and a *mini-white* gene as a selection marker. Each construct was integrated in the same attP landing site (ZH-51C), using phiC31-mediated recombination.

To test how well various transgenic Ash1 variants counteract Polycomb repression we turned to segment-specific expression of homeotic genes *Ubx* and *Abd-B* as a readout. *Ubx* and *Abd-B* are part of the bithorax complex gene cluster and specify the identity of the third thoracic and the first abdominal segments and the last four abdominal segments, respectively. (Lewis 1978). Both genes are classic targets of Polycomb/Trithorax regulation. When *ash1* function is compromised, the expression of *Ubx* and *Abd-B* is stochastically lost which, in turn, leads to partial transformation of corresponding segments towards the anterior fate (Shearn 1989; Klymenko and Muller 2004; Huang *et al*. 2017; Schmahling *et al*. 2018).

Of the published *ash1* alleles, *ash1^22^* and *ash1^Df(3L)Exel9011^* (hereafter referred to *ash1^9011^*) are the most severe. *ash1^22^* is a point mutation that converts Glutamin 129 into an early stop codon and has been reported as a null allele (Tripoulas *et al*. 1996). The *ash1^9011^* allele is the 53kb deletion that spans the entire *ash1* gene region as well as 13 other genes (Parks *et al*. 2004). *ash1*^22^/*ash1^9011^* animals die at early pupal stage with less than 10% of the animals producing enough adult cuticle to see homeotic transformations (Dorafshan *et al*. 2019). The hypomorphic *ash1^21^* allele is a substitution of Glutamic acid 1365 to Lysine within the Associated With SET (AWS) domain (Tripoulas *et al*. 1994). *ash1*^22^/*ash1^21^* animals develop to pharate adult stage and about 12% survive as adults (Dorafshan *et al*. 2019). The mutant adult flies show haltere to wing and third leg towards second leg transformations, reflecting partial loss of the *Ubx* gene expression (Figure S1A). They also show transformations of the 5^th^ and 6^th^ abdominal segments towards anterior fate caused by partial loss of the *AbdB* gene expression (Figure S1B).

To assess whether our transgenic system is capable to supply enough Ash1 protein, we introduced the transgene expressing full-length Ash1 (*Ubi-Ash1FL*) into the *ash1*^22^/*ash1^9011^* mutant background. Two copies of the *Ubi-Ash1FL* transgene fully restored the viability of *ash1*^22^/*ash1^9011^* flies (Figure S2), which showed no homeotic transformations, were fertile and could be maintained as a stock.

### Ash1 requires BAH, PHD and SET domains to counteract Polycomb repression

To test whether Ash1 variants lacking specific protein domains are able to counteract Polycomb repression, we crossed flies bearing one copy of a transgenic Ash1 variant on the second chromosome and the *ash1^9011^* allele on the third chromosome with flies containing one copy of the same transgenic Ash1 variant on the second chromosome and the *ash1*^22^ allele on the third chromosome (Figure 2A). To control for maternal effects, each cross was repeated in reciprocal setup. Four distinct classes are expected in the progeny of such cross (Figure 2B). If a transgenic Ash1 fully compensates for the loss of endogenous Ash1 protein, in the progeny, the number of trans-heterozygous *ash1*^22^/*ash1*^9011^ animals, supplemented with one or two copies of the Ash1 transgene should be one half of the number of flies heterozygous for *ash1* mutation (Figure 2B). We also expect transgenic *ash1*^22^/*ash1*^9011^ animals to have no homeotic transformations.

**Figure 2.**
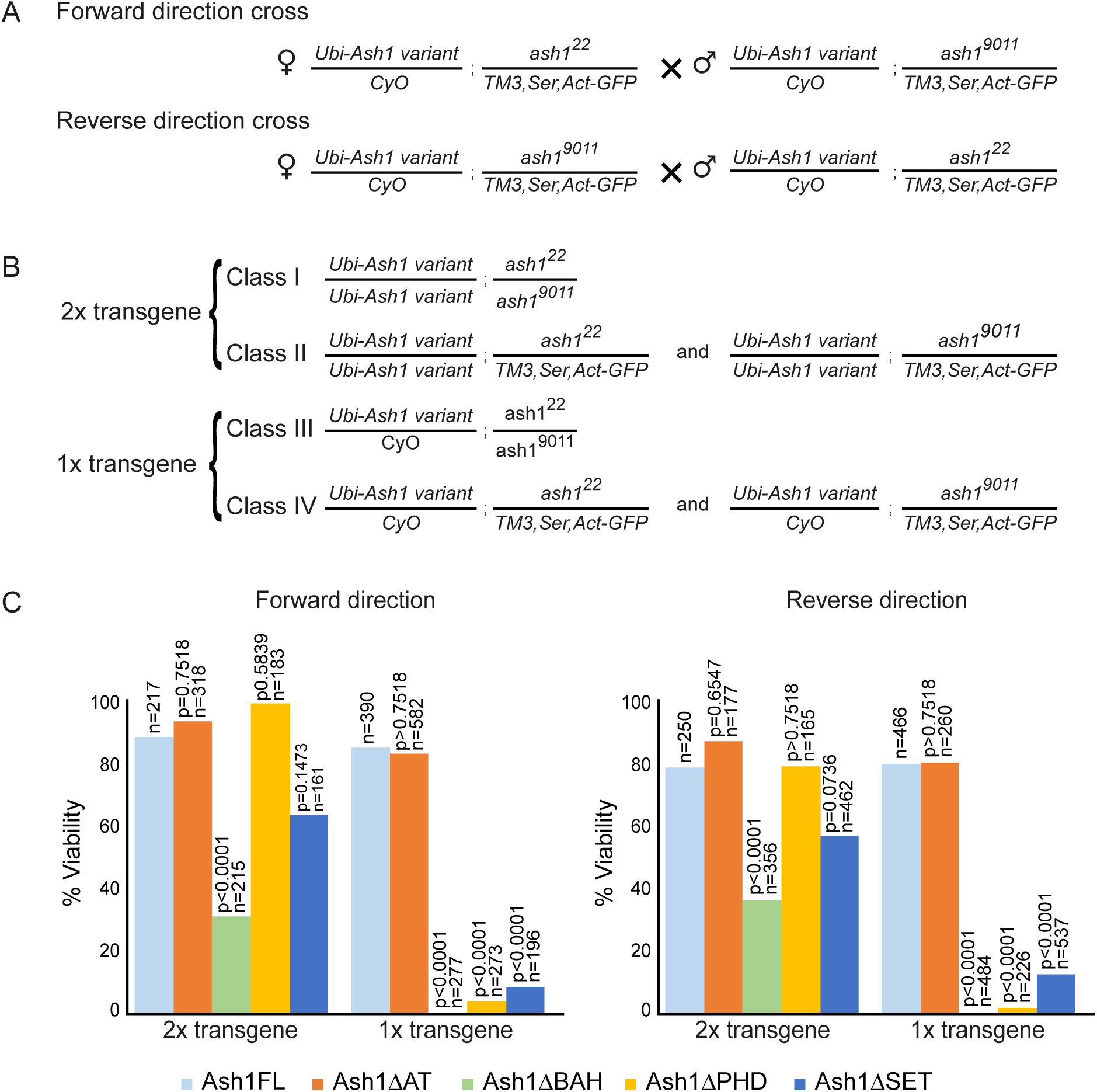
Complementation of *ash1^22^/ash1^9011^* mutations with transgenic Ash1 variants. **(A)** Reciprocal crossing schemes to combine different *Ash1* transgenes with the *ash1^22^/ash1^9011^* mutations. **(B)** The above crosses are expected to yield four phenotypically distinct progeny classes. Class I and Class II flies contain two copies of the Ash1 transgene while Class III and Class IV flies bear the marker *Cy* mutation and only one copy of the transgene. Class I and III flies have *ash1^22^/ash1^9011^* background while Class II and IV flies are heterozygous for only one of the mutant *ash1* alleles and carry *Ser* and *GFP* markers. **(C)** Histograms show the viability of *ash1^22^/ash1^9011^* mutant flies supplemented with different *Ash1* transgenes. The viability is plotted as percent ratios between the number of Class I and Class II progeny multiplied by two (for two copies of transgenic constructs) or as percent ratios between the number of Class III and Class IV progeny multiplied by two (for one copy of transgenic constructs). The ability of different *Ash1* mutant transgenes to restore the viability of *ash1^22^/ash1^9011^* flies was compared to that of the *Ash1FL* transgene and the significance of the deviation evaluated with *X*^2^ test (df=1). n = total number progeny counted for each cross.

Similar to *Ubi-Ash1FL*, one or two copies of the *Ash1* transgene lacking AT-hooks (*Ubi-Ash1ΔAT*) restored the viability and proper expression pattern of the homeotic genes (Figures 2C, 3A-B). In contrast, the transgenes lacking the BAH, PHD, or SET domains (*Ubi-Ash1ΔBAH*, *Ubi-Ash1ΔPHD* and *Ubi-Ash1ΔSET*) failed to complement *ash1*^22^/*ash1*^9011^ mutations. Although two copies of the transgenes could partially restore the viability of the *ash1*^22^/*ash1*^9011^ flies, one copy was not sufficient (Figure 2C). More importantly, *ash1*^22^/*ash1*^9011^ flies “rescued” with the two copies of ether transgene displayed obvious homeotic transformations (Figure 3A-B). Western-blot analysis showed that the *Ubi-Ash1ΔBAH*, *Ubi-Ash1ΔPHD* and *Ubi-Ash1ΔSET* transgenes produce at least as much protein as the transgene expressing the full-length Ash1 variant (Figure 3C-D). This rules out the trivial possibility that the Ash1 variants lacking BAH, PHD, or SET domains are less stable. Overall, we conclude that Ash1 requires BAH, PHD and SET domains, but not AT-hooks, to counteract Polycomb repression of homeotic genes.

**Figure 3.**
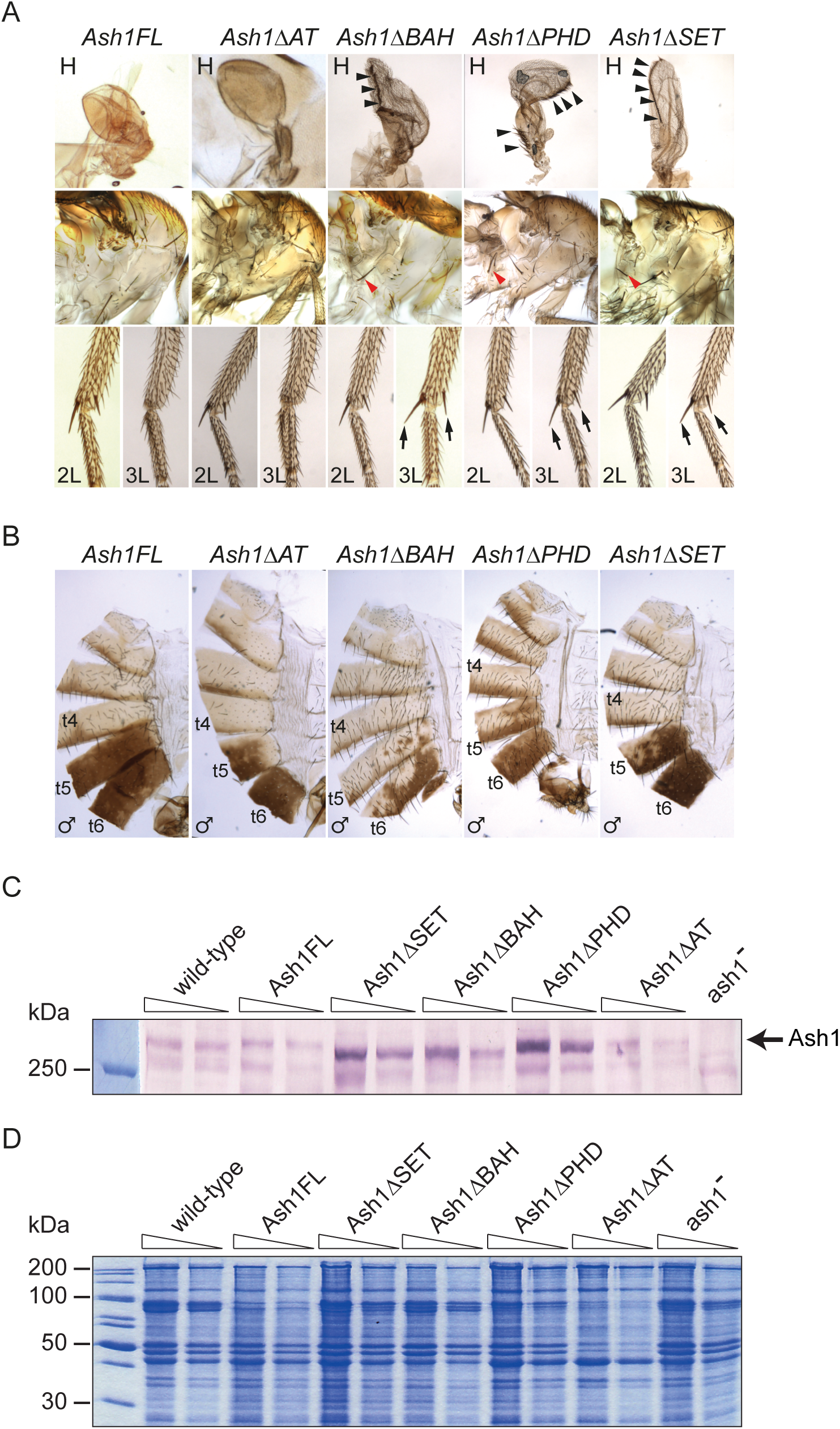
Homeotic phenotypes of *Ash1* transgenes. *ash1^22^/ash1^9011^* flies supplemented with *Ash1FL* or *Ash1ΔAT* transgenes appear wild type. This is in contrast to *Ash1ΔBAH*, *Ash1ΔPHD*, and *Ash1ΔSET* transgenes, which show transformations characteristic of erroneous repression of the *Ubx* and *Abd-B* genes. **(A)** Erroneous *Ubx* repression in the third thoracic segment (T3) and its partial transformation in the second thoracic segment is visible from transformation of haltere (H) to wing (note the change in shape and bristles marked with black arrowheads) the appearance of the hypopleural bristles on T3 (red arrowheads) and the presence of apical and pre-apical bristles on the third leg (3L) normally present only on the second leg (2L). (**B**) Erroneous repression of *Abd-B* and resulting transformation of posterior abdominal segments to more anterior fate is visible from partial loss of pigmentation in tergites 5 (t5) and 6 (t6). **(C)** Nuclear protein from the third instar larvae of wild-type animals, *ash1^22^/ash1^9011^* mutants (*ash1^-^*) and *ash1^22^/ash1^9011^* mutants supplemented with indicated *Ash1* transgenes was analyzed by western-blot with antibodies against Ash1. For each sample two 2-fold dilutions were loaded except for *ash1^22^/ash1^9011^* sample where only the least dilute material was used. Arrow indicates the position of Ash1. **(D)** Coomassie staining of the SDS-PAGE separated protein samples from (C) was used to control the loading.

### New alleles suggest that *ash1^22^* is a hypomorphic mutation

The observation that two copies of *Ubi-Ash1ΔBAH*, *Ubi-Ash1ΔPHD* and *Ubi-Ash1ΔSET* transgenes can restore the viability of *ash1^22^/ash1^9011^* flies was unexpected. Ash1 protein or mRNA deposited in the embryo by the heterozygous mother may help the mutant embryos to develop to adult stage. However, the *ash1^22^/ash1^9011^* flies supplemented with two copies of the *Ubi-Ash1ΔSET* or *Ubi-Ash1ΔPHD* transgenes are fertile and, although too weak to establish the permanent stock, can be interbred and propagated for up to three generations. This argues that maternal contribution is not the reason for their viability. Alternatively, Ash1 may have two distinct functions, one necessary to counteract Polycomb repression, another required for viability. Perhaps, the methyltransferase activity of the SET domain or the functions of the BAH and PHD domains are needed for the former but are dispensable for the viability. Finally, *ash1^22^* mutation may retain some wild-type function. This mutation introduces a premature stop codon at position 129 of the open reading frame, which truncates it to encode for a short polypeptide that lacks all conserved domains (Tripoulas *et al*. 1996). However, recent evidence indicates that, in many *Drosophila* mRNAs, ribosomes can read through single stop codon and produce small but functional amounts of polypeptides that incorporate amino acids encoded downstream of supposed translation termination sites. (Steneberg and Samakovlis 2001; Dunn *et al*. 2013). Therefore, the premature stop codon of the *ash1^22^* allele may be read through yielding small amount of the full-length protein, whose function may be boosted by large amounts of truncated Ash1 proteins.

To discriminate between the two possibilities, we generated new *ash1* alleles using CRISPR/Cas9 system (Gratz *et al*. 2014). To this effect, we designed guide RNAs to target Cas9 endonuclease to a site within the portion of *ash1* ORF that encodes the SET domain and to another site just downstream of the post-SET domain (Figure 4A). When cut at the two designated sites and repaired by the non-homologous end joining, the edited DNA would carry an ORF encoding for the catalytically inactive protein that lacks half of the SET domain and the entire post-SET domain (Figure 4A). Screening for such deletion, we recovered two new alleles. One corresponded to the desired “in-frame” deletion of 79 amino acids, which we called *ash1^7F^* (Figure 4A). Another deletion, dubbed *ash^3M^*, was larger (81 amino acids) and caused translation frame shift that introduced multiple successive stop codons in the middle of the SET domain (Figure 4A). The only possible product of the *ash^3M^* allele is a truncated Ash1 protein that lacks all but AT-hook domains and unlikely to be functional. Importantly, the heterozygous *ash^3M^ /+* flies are viable, fertile and show no homeotic transformations, which argues that *ash^3M^* is not a dominant negative mutation.

**Figure 4.**
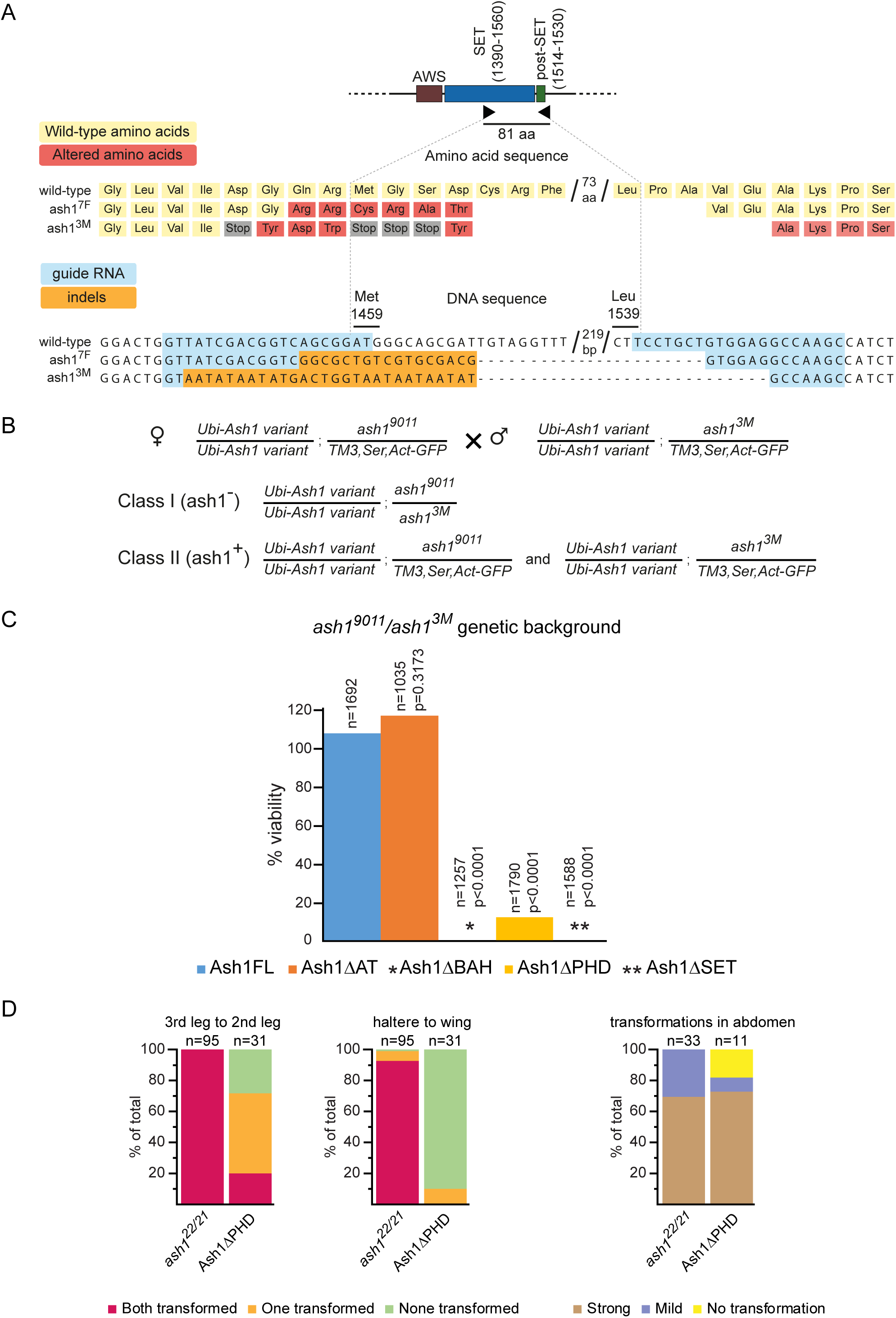
Generation and genetic analysis of new *ash1* alleles. **(A)** CRISPR/Cas9-mediated generation of new *ash1* alleles. Two guide RNAs (black triangles) were designed to generate dsDNA breaks at positions corresponding to amino acid 1459 (Met) and 1539 (Leu) within the ash1 ORF. In the screen for the deletion, two alleles were recovered. In the *ash1^7F^* allele 79 amino acid are deleted and 6 amino acids around the deletion breakpoint are changed (marked with red rectangles). In the *ash1^3M^* allele, 81 amino acid are deleted and a frameshift before the deletion breakpoint changed 4 amino acids (red rectangles) and introduced 4 stop codons (grey rectangles). The frameshift continues after the deletion breakpoint. In the schematic of the corresponding DNA sequences, the nucleotides highlighted in blue indicate positions of the guide RNAs and the nucleotides highlighted in orange represent insertions and deletions (indels). **(B)** Crossing scheme to test the complementation of *ash1^3M^/ash1^9011^* mutations with *ash1* transgenes. Female flies homozygous for an *ash1* transgene on the second chromosome and heterozygous for *ash1^9011^* allele on the third chromosome are crossed to males with the same second chromosome but heterozygous for *ash1^3M^* allele on the third chromosome. In the progeny, two classes are expected. While both classes are homozygous for an *ash1* transgene, Class I (*ash1^-^*) flies are trans-heterozygous for *ash1^3M^* and *ash1^9011^* alleles while Class II (*ash1^+^*) contains a wild-type *ash1* allele present on the balancer *TM3, Ser, Act-GFP* chromosome. The cross is expected to yield twice as many Class II flies. **(C)** The ability of truncated Ash1 proteins to restore the viability of *ash1^9011^/ash1^3M^* flies was compared to that of the full-length transgenic Ash1 (Ash1FL). While the *Ash1ΔAT* transgene fully restored the viability, neither *Ash1ΔBAH* nor *Ash1ΔSET* transgenes yielded any viable flies and the *Ash1ΔPHD* transgene complemented *ash1* loss of function inefficiently. n = the total number of progeny counted, p = probability that the observed difference in the survival rate of the wild-type (Ash1FL) and truncated *ash1* transgenes is caused by chance (as evaluated by *X*^2^ test, df=1) and % viability indicates the ClassI/ClassII ratio multiplied by two. **(D)** Homeotic phenotypes of the adult *ash1^22^/ash1^21^* (control) and *Ash1ΔPHD*; *ash1^9011^/ash1^3M^* flies. For representative images of strong and mild abdominal transformations see Figure S3. Y-axes display the percent of flies with corresponding transformation. n = the total number of flies scored.

Unlike *ash1^22^/ash1^9011^* flies supplemented with two copies of the *Ubi-Ash1ΔSET* transgene, homozygous *ash1^7F^* or trans-heterozygous *ash1^7F^/ash1^9011^* flies die as third instar larvae or at early pupal stage. This indicates that, when the possible source of the wild-type protein is excluded, the Ash1 SET domain is required for flies to survive. Which, in turn, discounts the idea that Ash1 has two separable functions, one necessary to counteract Polycomb repression and another for the viability. Attempts to complement the trans-heterozygous *ash1^9011^/ash1^3M^* mutations (likely zygotic null) with two copies of various *Ash1* transgenes further corroborate this notion (Figures 1A, 4B). In contrast to the *ash1^22^/ash1^9011^* complementation results (Figure 2C), the *Ubi-Ash1ΔBAH* and *Ubi-Ash1ΔSET* transgenes failed to restore the viability of the *ash1^9011^/ash1^3M^* mutants although the *Ubi-Ash1FL* and *Ubi-Ash1ΔAT* transgenes were still able to do so (Figure 4C). These observations argue that the *ash1^3M^* allele is more severe than *ash1^22^* and that the BAH, or SET domain-deficient Ash1 proteins have no intrinsic ability to support the fly viability. Consistent with the *ash1^3M^* allele being more severe, the *ash1^3M^* embryos show greater loss of the *Abd-B* expression in the central nervous system compared to their *ash1^22^/ash1^9011^* counterparts (Figure 5). Somewhat surprisingly, we noticed that about 10% of the *ash1^9011^/ash1^3M^* mutants supplemented with two copies of the *Ubi-Ash1ΔPHD* transgene survive to adult stage (Figure 4C). This is much smaller fraction compared to that detected in the *ash1^22^/ash1^9011^* complementation test (Figure 2C). Also, in contrast to the latter case, the survived *Ubi-Ash1ΔPHD/Ubi-Ash1ΔPHD; ash1^9011^/ash1^3M^* adults showed mild or no homeotic transformations (Figures 4D, Figure S3). Taken together, these results suggest that the PHD domain is needed for both, full fitness and the ability to counteract Polycomb repression, but is less critical compared to the SET and BAH domains. Overall, our results argue that *ash1^22^* is not a true loss-of-function allele but a strong hypomorph.

**Figure 5.**
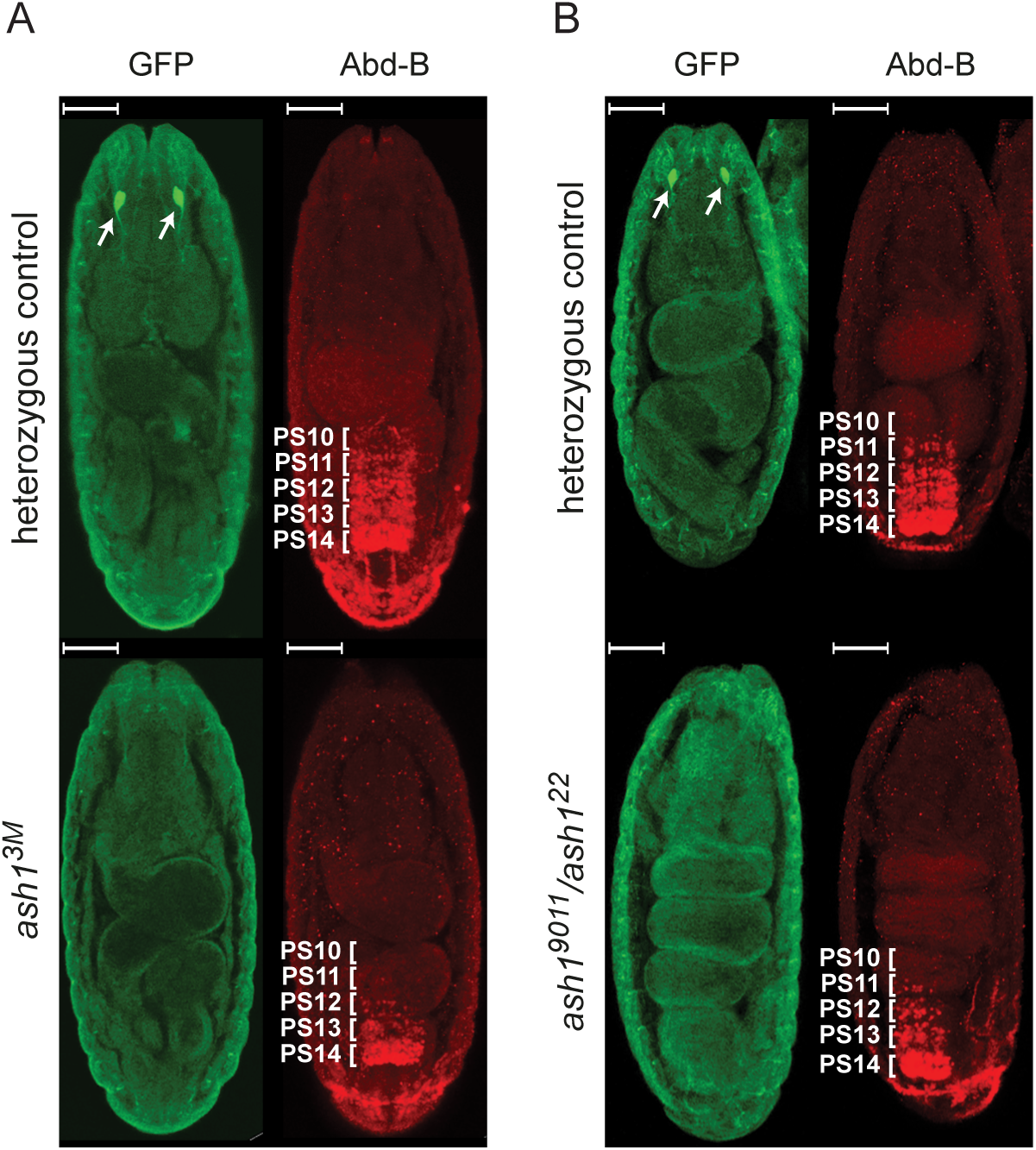
Stochastic loss of *Abd-B* expression in the central nervous system of the *ash1* mutant embryos. Immunostaining of *ash1* mutant embryos with antibodies specific to Abd-B (red) shows reduced Abd-B expression in both homozygous *ash1^3M^* (**A**) and trans-heterozygous *ash1^9011^/ash1^22^* (**B**) mutant embryos. Heterozygous embryos, where *ash1* mutant allele is combined with the *TM3,Sb,e,Kr::GFP* balancer, serve as a control. In the control embryos *Abd-B* is expressed in parasegments 14-10 (marked with white brackets) in a gradient that slopes towards anterior pole. The loss of Abd-B signal, although seen in both mutants, is more pronounced in homozygous *ash1^3M^* embryos compared to trans-heterozygous *ash1^9011^/ash1^22^* counterparts. In both (**A**) and (**B**), the control and mutant embryos were stained together and separated by strong GFP immunostaining (green) of the Bolwig’s organs (marked with white arrows). The embryos are oriented with anterior poles facing up. The scale bars correspond to 50 μm.

### Interallelic complementation indicates that Ash1 protein is modular and acts as a multimer

BAH, PHD and SET domains are all required for Ash1 to counteract Polycomb repression. While SET domain likely functions by methylating lysines (Tanaka *et al*. 2007; Huang *et al*. 2017; Schmahling *et al*. 2018) of yet unknown substrate(s) (Dorafshan *et al*. 2019), the roles of PHD and BAH domains are less clear. Genome-wide mapping in cultured *Drosophila* cells indicates that Ash1 binds weakly to multiple genomic sites, often located within long 5’ introns of transcriptionally active genes (Kharchenko *et al*. 2011; Huang *et al*. 2017). It also binds strongly to a few dozens of regions, many of which correspond to known Polycomb regulated genes caught in transcriptionally active state (Schwartz *et al*. 2010; Kharchenko *et al*. 2011; Huang *et al*. 2017). PHD and BAH domains may be involved in binding of Ash1 to either or both types of regions.

To investigate this possibility, we performed Chromatin Immunoprecipitation (ChIP) and immunostained polytene chromosomes with antibodies against Ash1. Ideally, we would have liked to examine the binding of various transgenic Ash1 variants in animals completely devoid of endogenous Ash1 protein (i.e. on *ash1^9011^/ash1^3M^* background). Unfortunately, such larvae proved too laborious to collect in numbers sufficient for ChIP. Therefore, we resorted to do the assays in *ash1^22^/ash1^9011^* mutants (strong hypomorphs). As illustrated by Figure 6, in the control *ash1^22^/ash1^9011^* larvae, ChIP signals and the polytene chromosome staining are reduced to background. However, both are restored to wild-type levels in larvae supplemented with two copies of transgenes expressing either the full-length or the PHD- or SET-deficient Ash1 proteins. This argues that the PHD- and SET-deficient proteins can still bind strongly to the de-repressed Polycomb regulated genes and weakly to multiple sites throughout the genome. Polytene chromosomes from *Ash1ΔBAH; ash1^9011^/ash1^22^* larvae show weaker anti-Ash1 staining suggesting that Ash1ΔBAH protein may bind chromatin less well compared to other tested variants. However, ChIP experiments did not detect obvious reduction in binding of the BAH-deficient protein so more work is needed to conclude whether BAH domain contributes to Ash1 binding to chromatin.

**Figure 6.**
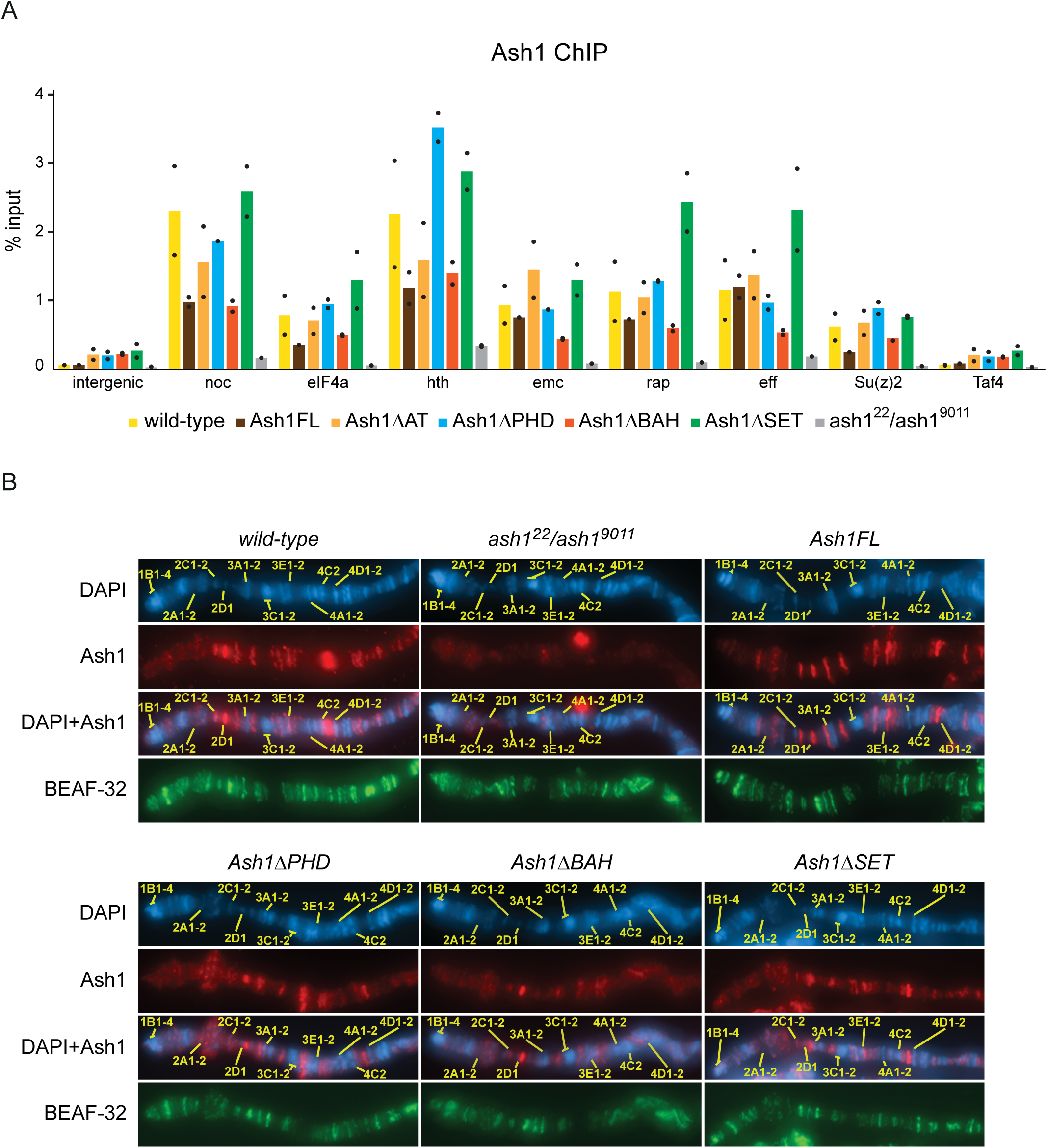
Truncated Ash1 proteins can still bind chromatin. (**A**) Chromatin from the wild-type, *ash1^9011^/ash1^22^*, *Ash1FL; ash1^9011^/ash1^22^* (Ash1FL), *Ash1ΔPHD; ash1^9011^/ash1^22^* (Ash1ΔPHD), *Ash1ΔBAH; ash1^9011^/ash1^22^* (Ash1ΔBAH), and *Ash1ΔSET; ash1^9011^/ash1^22^;* (Ash1ΔSET) third instar larvae was used for immunoprecipitation with the antibodies against Ash1. Histograms display the mean of the two independent experiments (n=2) with dots indicating individual experimental results. *noc*, *eIF4a*, *hth*, *emc*, *rap*, *eff* and *Su(z)2* represent loci strongly bound by Ash1 in wild-type larval cells. An intergenic region on chromosome 3R (intergenic) and constitutively expressed *Taf4* gene were used as negative controls. (**B**) Representative pictures of the distal part of the polytene X chromosome from the third instar larvae of the same genotypes as in (**A**) immunostained with antibodies against Ash1 protein (red). Immunostaining with antibodies against an unrelated BEAF-32 protein (green) was used as positive control. Staining with DAPI (blue) was used to map positions of characteristic polytene chromosome bands (yellow lines) according to Bridges nomenclature (Bridges 1935). No distinct bands are visible on the chromosomes of the *ash1^9011^/ash1^22^* mutants stained with anti-Ash1 antibodies, but the immunostaining pattern is restored in larvae supplemented with *ash1* transgenes. The chromosomes from *Ash1ΔBAH; ash1^9011^/ash1^22^* larvae show weaker anti-Ash1 staining suggesting that Ash1ΔBAH protein may bind chromatin less well compared to other tested variants.

To gain further insight into the contribution of individual domains, we considered two possible scenarios. In the first scenario, the SET, PHD and BAH domains work in concert as parts of the same Ash1 molecule. Possibly, enabling one specific molecular property, for example, efficient lysine methylation by the SET domain. Alternatively, BAH, PHD and SET domains may act as independent modules each contributing its own distinct function. If Ash1 molecules form multimers (or at least a dimer), the second scenario allows for the functional Ash1 protein to be composed from a mixture of distinct individually truncated Ash1 polypeptides. The latter could be genetically tested by interallelic complementation. In such cases, highlighted by classical experiments with *Neurospora crassa* glutamate dehydrogenase (Coddington and Fincham 1965) and bacteriophage T4 DNA Polymerase (Reha-Krantz 1990), the two alleles of the same gene encoding products defective at distinct protein domains may produce no function when expressed individually in the cell. However, when co-expressed in the same cell, their products physically interact, which restores their function.

To test whether, *in vivo*, Ash1 acts as a multimer and BAH, PHD and SET domains correspond to separable molecular modules, we performed two sets of complementation tests. First, we introduced different *ash1* transgenes into *ash1^7F^* mutant background. *ash1^7F^* allele has part of the SET domain and the entire post-SET domain deleted and encodes the protein that lacks metyltransferase activity but has all other domains intact. Strikingly, two copies of the control *Ubi-Ash1FL* as well as *BAH*- and *PHD*-deficient transgenes, but not the *Ubi-Ash1ΔSET* transgene, restore the viability of *ash1^7F^* mutant flies (Figure 7). These results argue that the BAH and PHD domains present on the SET-deficient Ash1 molecule produced by the *ash1^7F^* allele can combine their functions with that of the SET domain from a different molecule supplied in *trans-*. To test this further, we attempted to restore the viability of the *ash1^9011^/ash1^3M^* loss-of-function mutants, by expressing different combinations of truncated Ash1 proteins from two different transgenic constructs. As illustrated by Figure 8, the combinations of *Ash1ΔPHD/Ash1ΔSET* and *Ash1ΔBAH/Ash1ΔSET* transgenes partially restored the viability of the *ash1* loss of function mutants, while the combination of the *Ash1ΔPHD/Ash1ΔBAH* transgenes failed to do so. Some of the “rescued” *Ash1ΔPHD/Ash1ΔSET; ash1^9011^/ash1^3M^* flies appeared completely wild-type and others had only mild homeotic transformations (Figure 8C). Taken together, the results of interallelic complementation experiments argue that, *in vivo*, Ash1 functions as a multimer and that its SET domain acts as a distinct module that can be combined with BAH and PHD domains from different Ash1 peptide (Figure 9).

**Figure 7.**
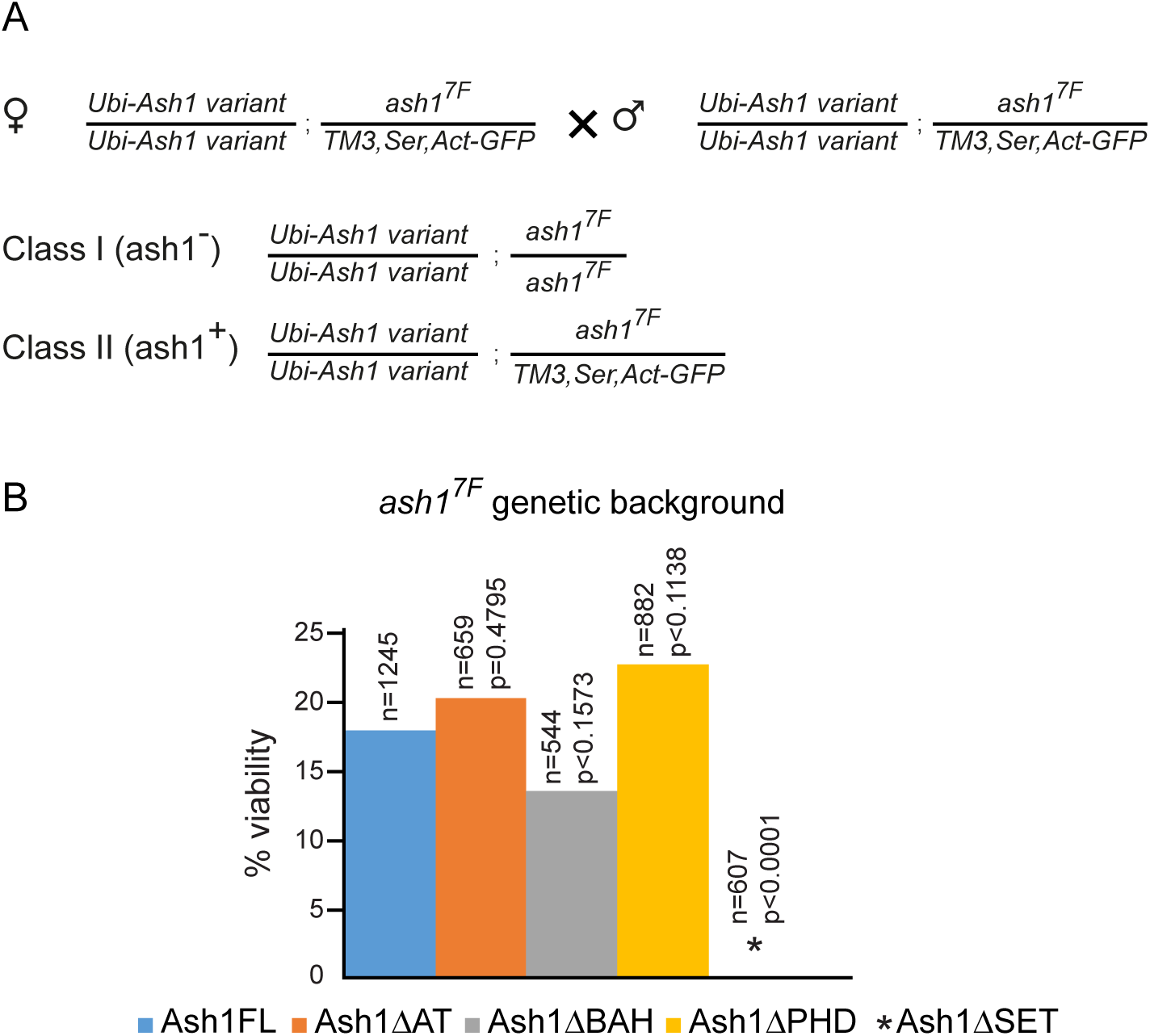
PHD- and BAH-deficient Ash1 proteins complement *ash1^7F^* mutation when provided in *trans*. **(A)** Crossing scheme to test the complementation of *ash1^7F^* mutation with *ash1* transgenes. Female flies homozygous for an *ash1* transgene on the second chromosome and heterozygous for *ash1^7F^* allele on the third chromosome were crossed to males with the same genotype. In the progeny, two classes are expected. While both classes are homozygous for an *ash1* transgene, Class I flies are homozygous for *ash1^7F^* mutation (*ash1^-^*) while Class II contains a wild-type *ash1* allele (*ash1^+^*). The cross is expected to yield twice as many Class II flies. **(B)** The ability of truncated Ash1 proteins to restore the viability of *ash1^7F^* mutants was compared to that of the full-length transgenic Ash1 (Ash1FL). While the *Ash1ΔAT*, *Ash1ΔBAH* and *Ash1ΔPHD* transgenes restore the viability to the same extent as *Ash1FL*, the *Ash1ΔSET* transgene does not. n = the total number of progeny counted, p = probability that the observed difference in the survival rate of the wild-type (*Ash1FL*) and truncated *ash1* transgenes is caused by chance (as evaluated by *X*^2^ test) and % viability indicates the ClassI/ClassII ratio multiplied by two. Note that even the viability of the *Ash1FL; ash1^7F^* flies remains at ∼20% of the expected, which suggests that the *ash1^7F^* chromosome carries additional unrelated recessive mutations that affect fly fitness.

**Figure 8.**
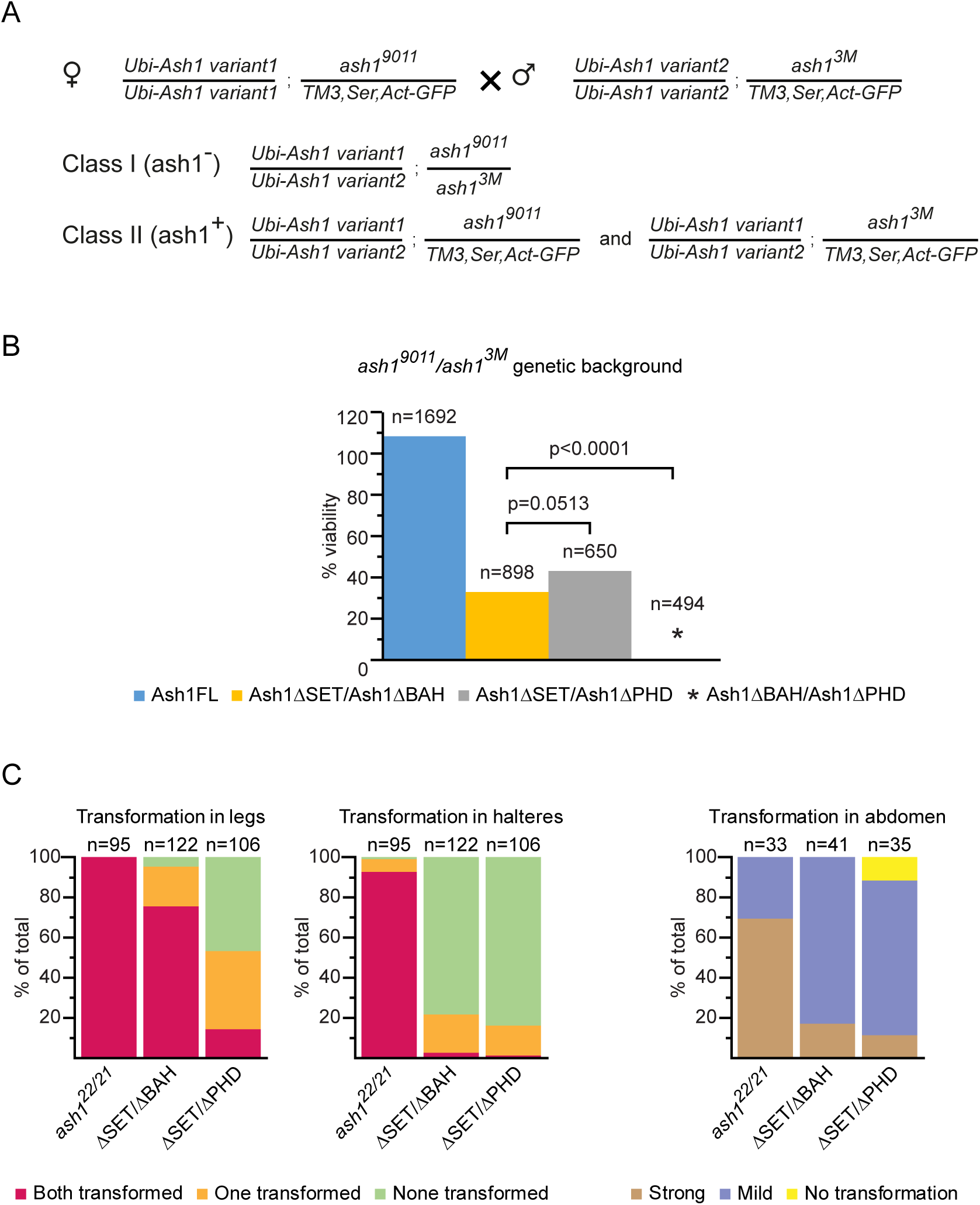
Truncated Ash1 proteins combine to partially restore the viability of *ash1* loss of function mutants. **(A)** Crossing scheme to test the complementation of *ash1^3M^/ash1^9011^* mutations with different combinations of *ash1* transgenes. Flies homozygous for one of the *ash1* transgenes on the second chromosome and heterozygous for *ash1^9011^* allele, balanced over *TM3,Ser,Act-GFP* third chromosome, were crossed with flies homozygous for another *ash1* transgene and heterozygous for the *ash1^3M^* allele, balanced over *TM3,Ser,Act-GFP* third chromosome. The same crossing scheme was used for all combinations of *Ash1ΔBAH*, *Ash1ΔPHD* and *Ash1ΔSET* transgenes. In the progeny, two classes are expected. While both classes are trans-heterozygous for a combination of the *ash1* transgenes, Class I flies are also trans-heterozygous for *ash1^3M^* and *ash1^9011^* alleles (*ash1^-^*) while Class II contains a copy of the wild-type *ash1* allele (*ash1^+^*). The cross is expected to yield twice as many Class II flies. **(B)** Different combinations of truncated ash1 transgenes were compared for their ability to complement the lethality of the *ash1^9011^*/ *ash1^3M^* mutants. The complementation by two copies of the *Ash1FL* transgene was used as positive control. n = the total number of progeny counted, p = probability that the observed difference in the survival rate of the *ash1* mutant flies supplemented with corresponding combinations of *ash1* transgenes is caused by chance (as evaluated by *X*^2^ test). % viability indicates the ClassI/ClassII ratio multiplied by two. **(C)** Quantification of homeotic transformations in the *ash1^9011^/ash1^3M^* flies supplemented with different combinations of *ash1* transgenes. Third to second leg, haltere to wing and male abdominal tergite 5 (t5) to tergite 4 (t4) transformations were evaluated in the Class I adult progeny of the cross in (**A**). Y-axis show the percent of flies with indicated transformation. n = the number of flies counted. Phenotypes of the *ash1^22^/ash1^21^* (*ash1^22/21^*) mutant flies were quantified as a benchmark. Representative images for the range of haltere to wing transformations as well as the strong and mild t5 to t4 transformation are shown in Figure S3.

**Figure 9.**
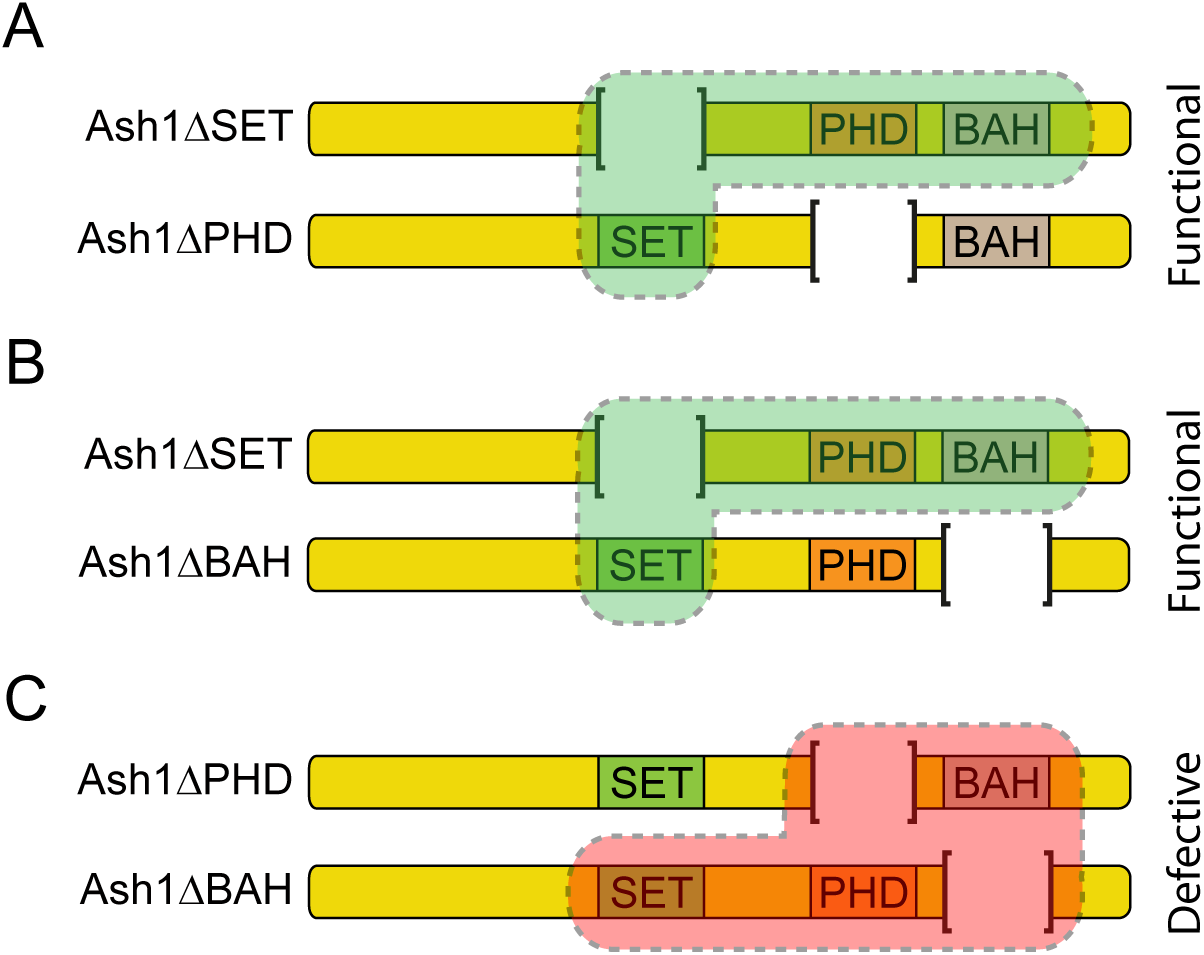
*Trans-* complementation between Ash1 domains. *In vivo*, Ash1 molecules form dimers (as shown on the schematics) or multimers. The SET domain of Ash1 can function as an autonomous module. Therefore, Ash1 molecules that lack the SET domain (Ash1ΔSET) can combine with molecules that lack BAH- or PHD-domains to yield functional Ash1 complex **(A, B)**. In contrast, a complex composed of the Ash1 molecules lacking either BAH- or PHD-domain appears ineffectual **(C)**. This suggests that BAH and PHD domains need to be part of the same molecule to function properly or may need to cooperate to allow multimerization.

## Discussion

This study leads to three main conclusions. First, we have shown that, in addition to the methyltransferase activity of the SET domain, the functions of the PHD and BAH domains are necessary for Ash1 to counteract Polycomb repression and support fly viability. Unexpectedly, evolutionarily conserved AT-hook domains appear not critical for either, although, we cannot exclude that they contribute to some aspect of Ash1 function not detected by our experiments. Our findings urge more effort to understand molecular properties of the Ash1 PHD and BAH domains. In other chromatin regulators both kinds of domains are known to interact with N-terminal tails of histones H3 or H4. Depending on variations in their amino acid sequences, PHD and BAH domains may have higher affinities to tails methylated at specific Lysines or Arginines (Sanchez and Zhou 2011; Yang and Xu 2013). In addition, some of the PHD domains can bind DNA or RNA (with dissociation constants comparable to those for interactions with histones) (Weaver *et al*. 2018) and many BAH domains mediate interactions with non-histone proteins (Yang and Xu 2013). Curiously, it was recently reported that BAH domains of the *Arabidopsis thaliana* proteins SHORT LIFE (SHL) and EARLY BOLTING IN SHORT DAYS (EBS) can specifically interact with the histone H3 tail methylated at Lysine 27 (Li *et al*. 2018). Should the BAH domain of Ash1 possess similar property, it would give Ash1 a neat way to discriminate between “regular” active genes and genes that just overcame Polycomb repression and still retain its hallmark, the histone H3 tri-methylated Lysine 27 (Schwartz *et al*. 2010).

Second, from the results of the interallelic complementation experiments, we conclude that the SET domain of Ash1 and the combination of its PHD and BAH domains (PHD-BAH cassette) represent two molecularly separable modules. This implies that many biochemical properties of the PHD-BAH cassette can be studied in isolation from the SET domain using shorter truncated Ash1 peptides. This is particularly advantageous given technical difficulties of working with the full-length protein of Ash1 size (over 250kDa). Our complementation experiments suggest that the PHD and BAH domains need to be on the same molecule to function properly. Yet, we note that small fraction of *ash1^9011^/ash1^3M^* flies survive to adult stage when supplemented with two copies of *Ubi-Ash1ΔPHD* transgene. (Figures 4C). The survival rate is poor, just 10% of that seen with the transgene expressing full-length Ash1. Nevertheless, this observation suggests that, while the PHD and BAH domains act in cooperative manner, the latter is still partially functional even in the absence of the PHD domain.

The third conclusion from this study is that *in vivo* Ash1 acts as a multimer. Additional biochemical studies are required to define the exact Ash1 tertiary structure. Regardless, the multimeric nature of the Ash1 protein is consistent with the dose dependent behavior of *ash1* mutations (Shearn 1989; Tripoulas *et al*. 1994). Moreover, it provides a neat explanation of why the function of the small amount of wild-type protein produced by the hypomorphic *ash1^22^* allele is boosted by defective Ash1 molecules produced from the transgenic constructs. Suppose Ash1 acts as a dimer. The equilibrium concentration of the homodimer is proportional to the square of the concentration of its constituent monomers. Hence, in the *ash1^22^* mutants, which produce only small amount of the full-length Ash1 peptides, the concentration of the dimers is going to be negligible. When large amounts of truncated Ash1 polypeptides are introduced, the concentration of the dimers incorporating at least one full-length Ash1 polypeptide becomes proportional to the concentration of the full-length polypeptides produced by the *ash1^22^* allele. Although still low compared to that in the wild-type cells, such dimer concentration is much higher than that in the *ash1^22^* mutants. The rationale behind Ash1 multimerization is currently unknown and needs further investigation. It is tempting to speculate that the ability to multimerize may help Ash1 to form broad chromatin domains over the de-repressed Polycomb-regulated genes (Schwartz *et al*. 2010; Kharchenko *et al*. 2011; Huang *et al*. 2017).

It has been commonly assumed that the early premature stop codon of the *ash1^22^* allele completely abolishes *ash1* function. Therefore, when it was found that the stochastic loss of the homeotic gene expression is detected only in *ash1^22^* homozygous embryos derived from the *ash1^22^* mutant germ cells, this was taken to indicate that maternally supplied Ash1 protein is sufficient to maintain the expression of homeotic genes throughout embryonic development. Since *ash1^22^* mutant animals derived from *ash1^22^* mutant germ cells survived until late larval stage, it seemed that the *ash1* function was less critical than that of the *trithorax* (*trx*) gene whose mutants die during embryogenesis. Our observations argue that the lack of detectable changes in the homeotic gene expression in the *ash1^22^* embryos produced by the heterozygous mothers is due to small amount of function provided by the *ash1^22^* allele and that such changes are readily detectable in mutants with combination of stronger *ash1* alleles. This, in turn, suggests that the phenotype of the true maternal and zygotic loss of *ash1* function is still unknown and that the question of whether *trithorax* and *ash1* functions are equally important is still open.

## Acknowledgements

We thank Dr. Maria Kim and Dr. Jan Larsson for gifts of fly strains and plasmids. We are grateful to Dr. Vincenzo Pirrotta for the gift of anti-Ash1 antibody. This work was supported in part by grants from Swedish Research Council to YBS and grants from Knut and Alice Wallenberg Foundation and Kempestiftelserna to EpiCoN (YBS co-PI). Stocks obtained from the Bloomington Drosophila Stock Center (NIH P40OD018537) were used in this study. The anti-Abd-B 1A2E9, monoclonal antibody developed by S. Celniker and the anti-BEAF-32 monoclonal antibody developed by P. Schedl were obtained from the Developmental Studies Hybridoma Bank, created by the NICHD of the NIH and maintained at the University of Iowa, Department of Biology, Iowa City, IA 52242.

## Author contributions

ED and YBS conceived the project. ED performed all genetic experiments and analyzed homeotic phenotypes and embryonic *Abd-B* expression. TGK performed ChIP experiments and characterized the expression of *Ash1* transgenes. AG analyzed Ash1 binding to polytene chromosomes. MS helped to analyze homeotic phenotypes. ED and YBS wrote the manuscript with input from all authors.

## Figure legends

**Figure S1.**
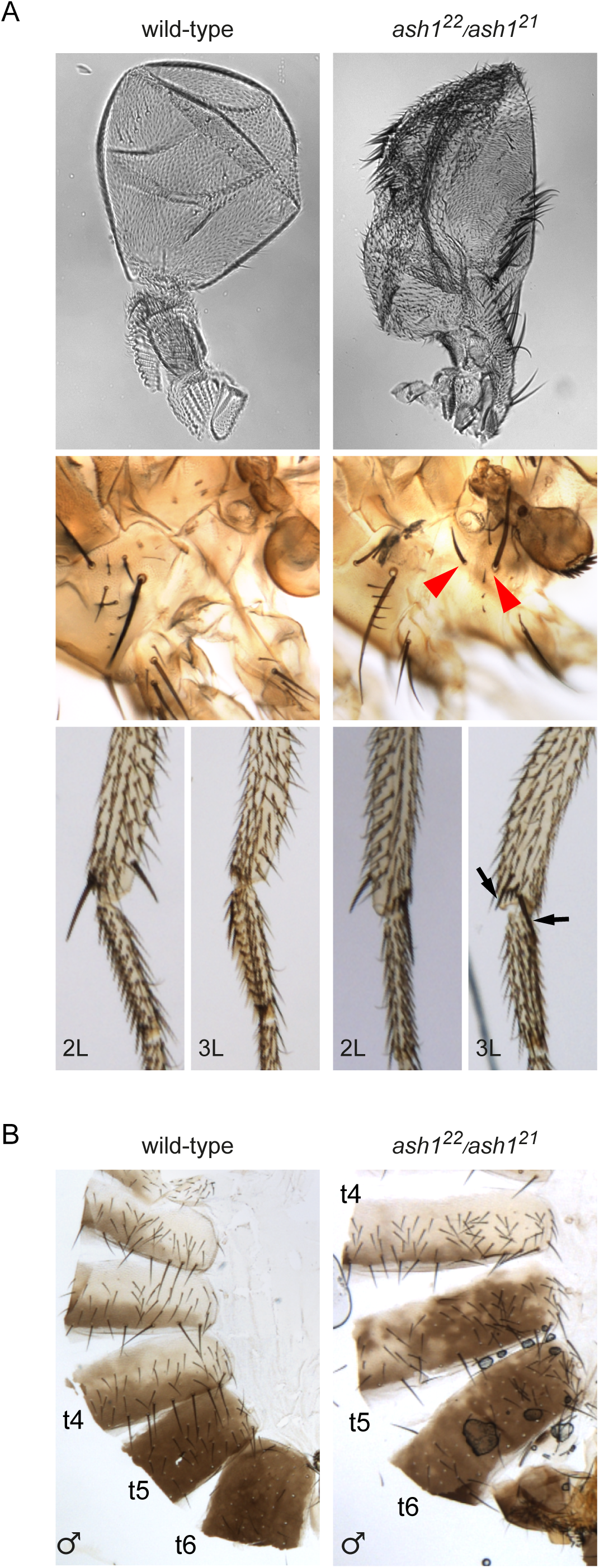
Homeotic transformation in *ash1* mutant flies. Characteristic homeotic phenotypes of *ash1* loss of function are caused by ectopic repression of *Ubx* and *Abd-B* genes. **(A)** Loss of *Ubx* expression leads to homeotic transformation of the third thoracic (T3) to second thoracic (T2) segment. This includes partial transformation of haltere to wing, characterized by the appearance of bristles on the haltere, appearance of hypopleural bristles on T3 (red arrowheads), and the third leg (3L) to second leg (2L) transformation manifested by the appearance of apical and pre-apical bristles on the third leg of *ash1^22^/ash1^21^* animals (black arrows). (**B**) Erroneous repression of *Abd-B* leads to transformation of posterior abdominal segments to more anterior fate. This is visible by the partial loss of pigmentation in tergites 5 (t5) and 6 (t6).

**Figure S2.**
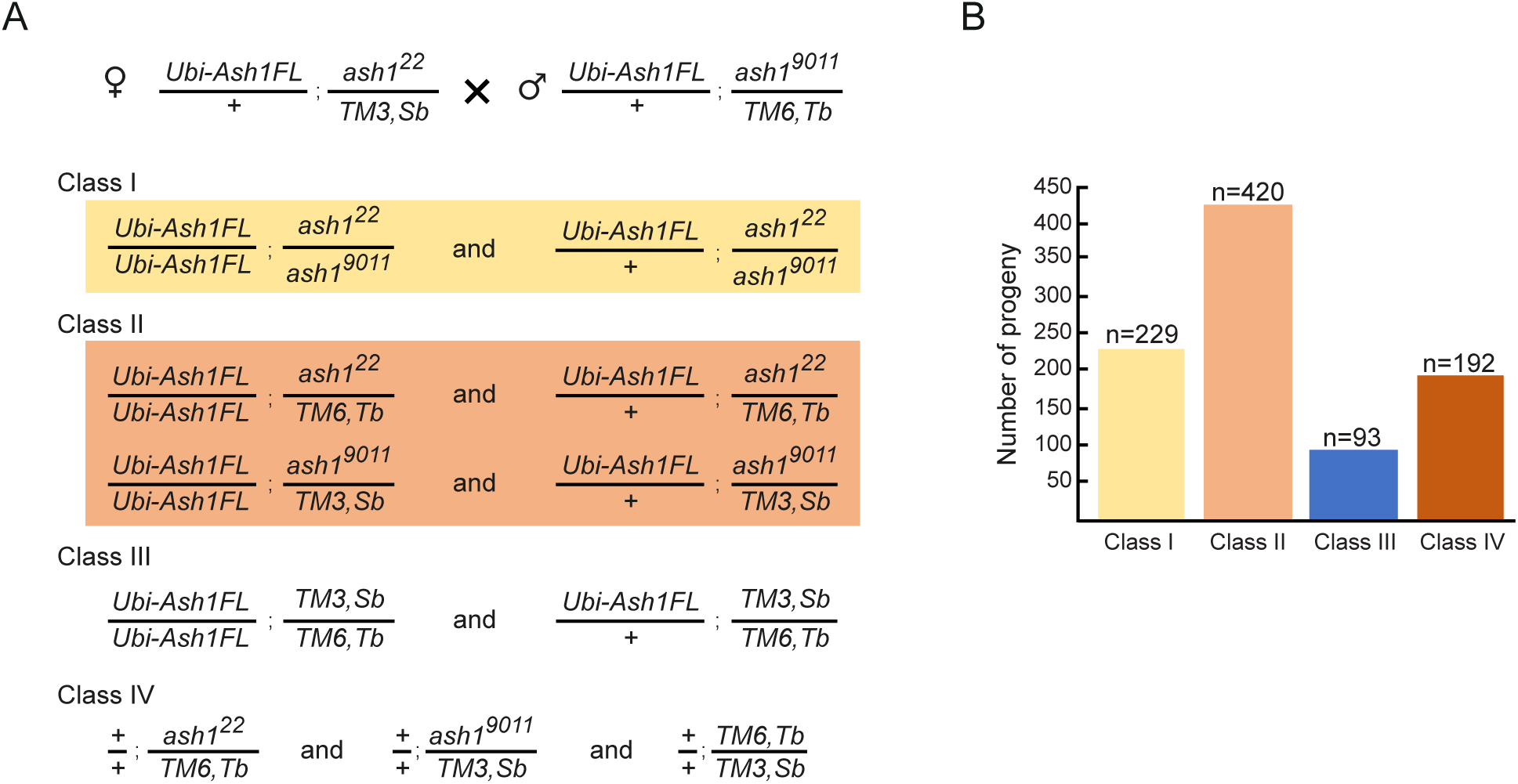
Expression from the *Ubi-p63E* promoter provides enough transgenic Ash1 protein. **(A)** Crossing scheme. Flies heterozygous for the full-length *Ash1* transgene under the control of *Ubi-p63E* promoter (*Ubi-Ash1FL*) on the second chromosome and heterozygous for the *ash1^22^* allele on the third chromosome were crossed to flies heterozygous for the same *Ash1* transgene and *ash1^9011^* allele. **(B)** The viability test. If the *Ash1* transgene fully complements the lethality of the *ash1^22^/ash1^9011^* mutants, the number of the Class I progeny is expected to be half of that of Class II. Lower than expected numbers of flies in Class III and Class IV are due to reduced fitness of flies with balancer chromosomes. (n = number of flies counted per phenotypic class).

**Figure S3.**
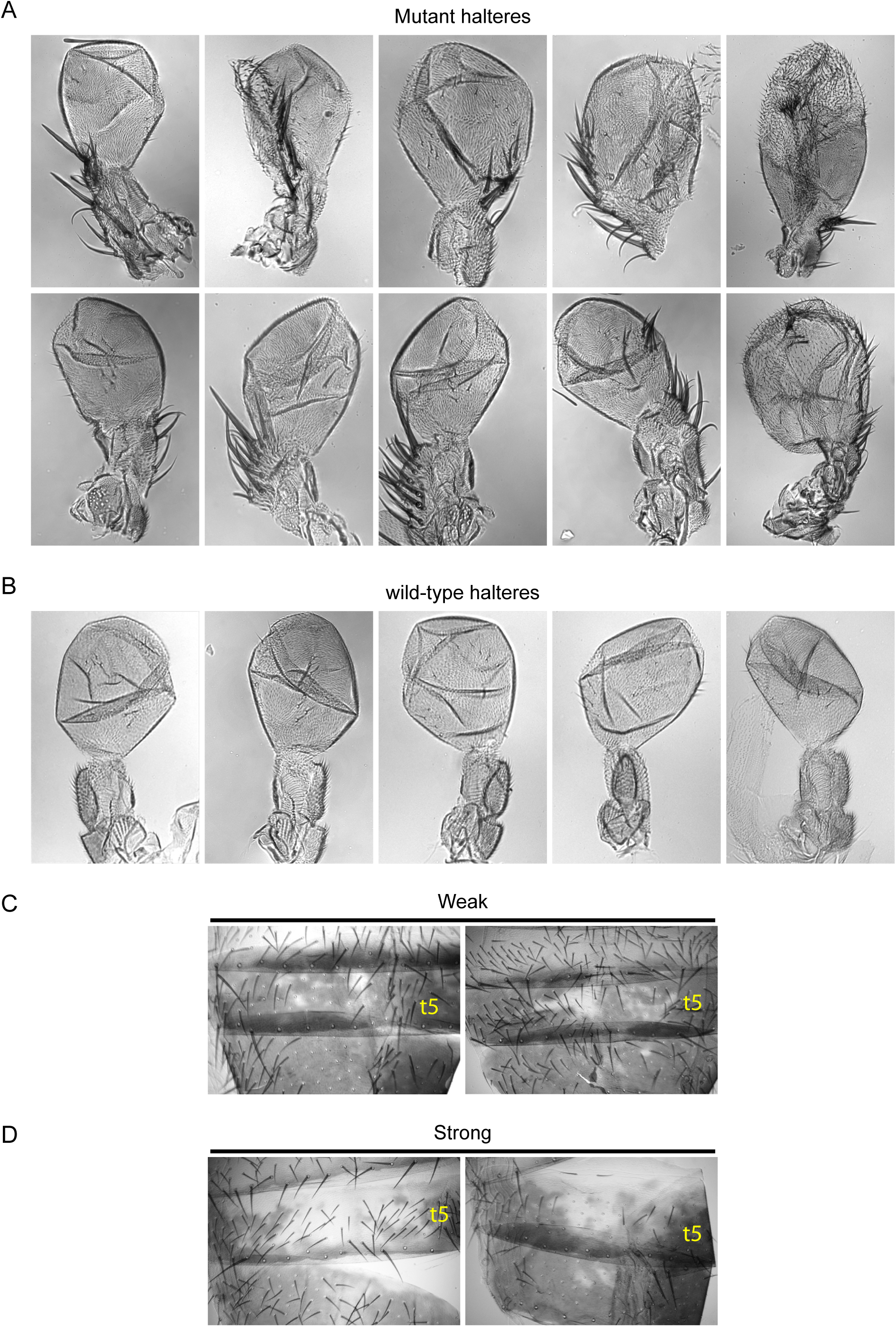
Representative pictures of wild-type and transformed halteres and male abdomens from *ash1* mutant flies. **(A)** Partial haltere to wing transformations ranging from mild (upper left corner) to severe (lower right corner). Note the appearance of bristles and changes in haltere shapes. (**B**) Wild-type halteres as a control. (**C**) Weak transformation of the abdomen in male flies is manifested by partial loss of pigmentation in tergite 5 (t5). (**D**) Strong transformation of the abdomen is characterized by greater area of t5 that lacks pigmentation.

**Table S1.**
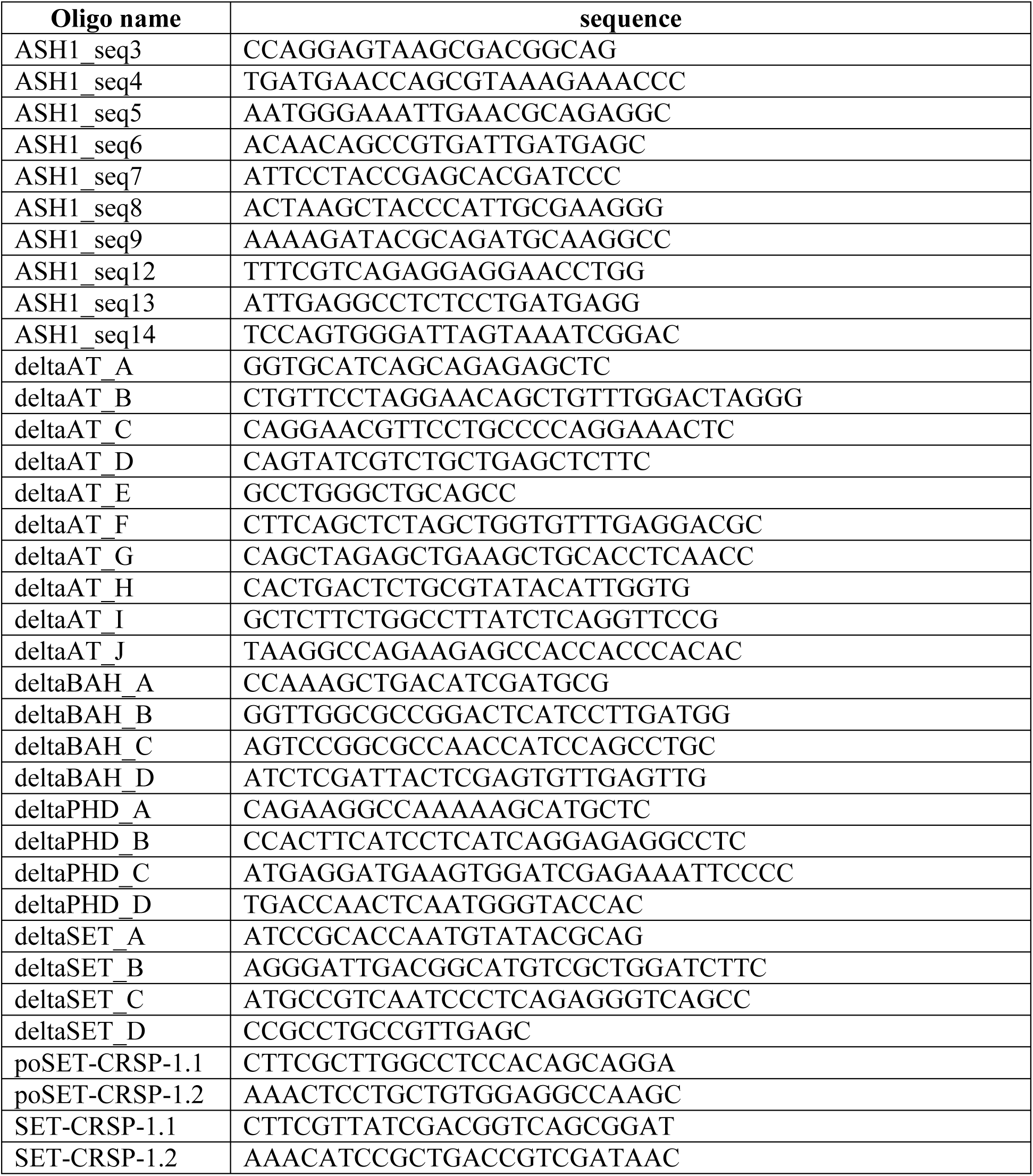
The list of oligonucleotides.

**Table S2.** The list of antibodies.

| **Antigen** | **Host** | **Reference/company/catalogue #** | **IF** | **ChIP** | **WB** |
| --- | --- | --- | --- | --- | --- |
| Abd-B | mouse monoclonal | DHSB, 1A2E9 | 1:10 | - | - |
| GFP | mouse monoclonal | Thermofisher, MA5-15256 | 1:200 | - | - |
| BEAF-32 | mouse monoclonal | DHSB, anti-BEAF | 1:5 | - | - |
| Ash1 | rabbit polyclonal | Kahn at al., 2016 | 1:500 | 1:100 | - |
| Ash1 | rabbit polyclonal | against aa 2156-2217, gift from V.Pirrotta | - | - | 1:500 |
| anti-rabbit Alexa-488 | goat | Invitrogen, # A-11008 | 1:500 | - | - |
| anti-mouse Alexa-555 | goat | Abcam, # ab150114 | 1:500 | - | - |
| anti-rabbit (Fc) AP conjugated | goat | Promega, #S3731 | - | - | 1:10000 |

**Table S3.**
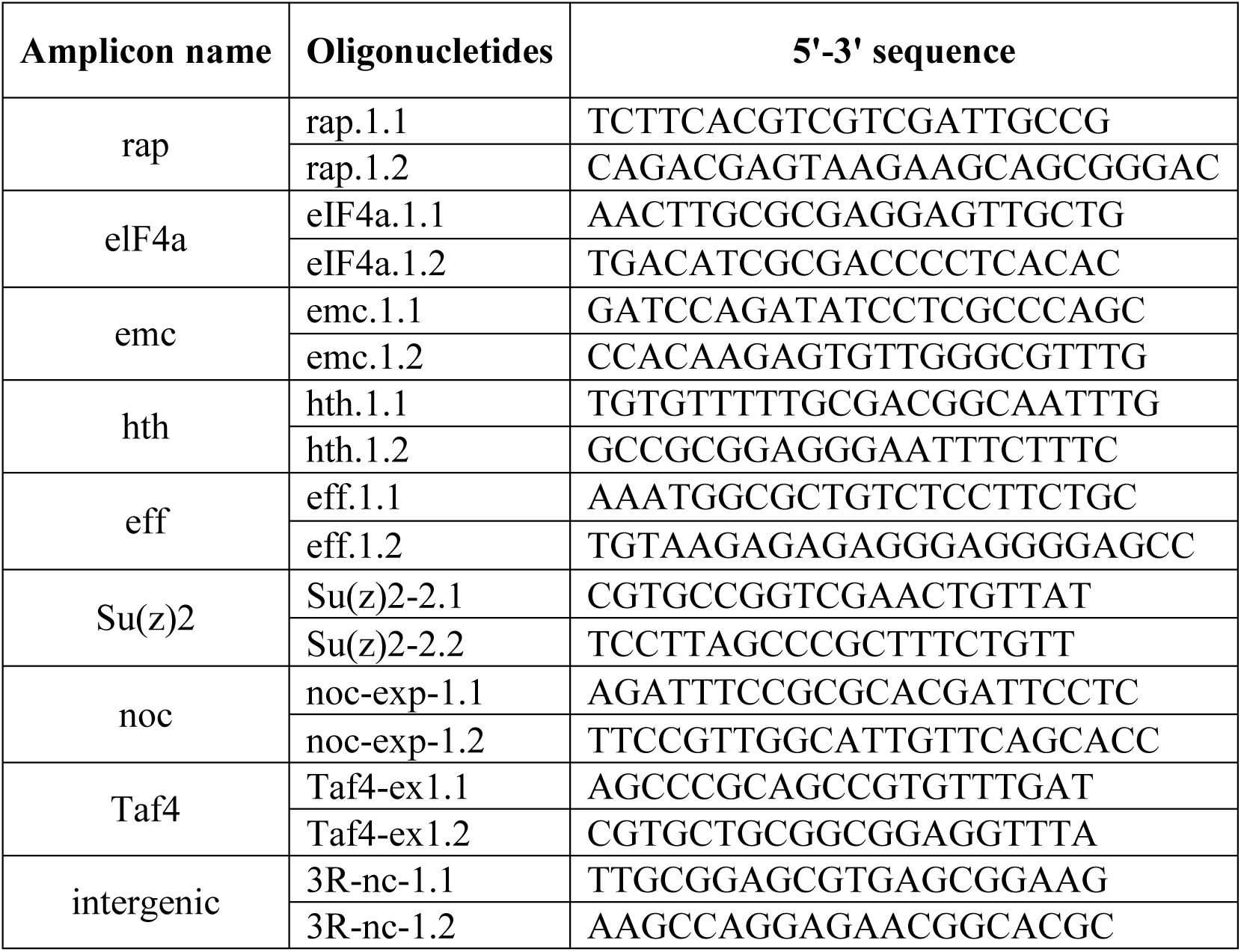
The list of amplicons used in ChIP-qPCR.

## References

An, S., K. J. Yeo, Y. H. Jeon and J. J. Song, 2011 Crystal structure of the human histone methyltransferase ASH1L catalytic domain and its implications for the regulatory mechanism. J Biol Chem 286: 8369–8374.

Bischof, J., R. K. Maeda, M. Hediger, F. Karch and K. Basler, 2007 An optimized transgenesis system for Drosophila using germ-line-specific phiC31 integrases. Proc Natl Acad Sci U S A 104: 3312–3317.

Bridges, C. B., 1935 Salivary chromosome maps with a key to the banding of the chromosomes of Drosophila melanogaster. J. Hered 26: 60–64.

Butcher, R. D., S. Chodagam, R. Basto, J. G. Wakefield, D. S. Henderson et al., 2004 The Drosophila centrosome-associated protein CP190 is essential for viability but not for cell division. J Cell Sci 117: 1191–1199.

Coddington, A., and J. R. Fincham, 1965 Proof of Hybrid Enzyme Formation in a Case of Inter-Allelic Complementation in Neurospora Crassa. J Mol Biol 12: 152–161.

Coleman, R. T., and G. Struhl, 2017 Causal role for inheritance of H3K27me3 in maintaining the OFF state of a Drosophila HOX gene. Science 356.

Dorafshan, E., T. G. Kahn, A. Glotov, M. Savitsky, M. Walther et al., 2019 Ash1 counteracts Polycomb repression independent of histone H3 lysine 36 methylation. EMBO Rep 20.

Dorighi, K. M., and J. W. Tamkun, 2013 The trithorax group proteins Kismet and ASH1 promote H3K36 dimethylation to counteract Polycomb group repression in Drosophila. Development 140: 4182–4192.

Dunn, J. G., C. K. Foo, N. G. Belletier, E. R. Gavis and J. S. Weissman, 2013 Ribosome profiling reveals pervasive and regulated stop codon readthrough in Drosophila melanogaster. Elife 2: e01179.

Eggert, H., A. Gortchakov and H. Saumweber, 2004 Identification of the Drosophila interband-specific protein Z4 as a DNA-binding zinc-finger protein determining chromosomal structure. J Cell Sci 117: 4253–4264.

Gratz, S. J., F. P. Ukken, C. D. Rubinstein, G. Thiede, L. K. Donohue et al., 2014 Highly specific and efficient CRISPR/Cas9-catalyzed homology-directed repair in Drosophila. Genetics 196: 961–971.

Hodl, M., and K. Basler, 2012 Transcription in the absence of histone H3.2 and H3K4 methylation. Curr Biol 22: 2253–2257.

Hou, P., C. Huang, C. P. Liu, N. Yang, T. Yu et al., 2019 Structural Insights into Stimulation of Ash1L’s H3K36 Methyltransferase Activity through Mrg15 Binding. Structure 27: 837–845 e833.

Huang, C., F. Yang, Z. Zhang, J. Zhang, G. Cai et al., 2017 Mrg15 stimulates Ash1 H3K36 methyltransferase activity and facilitates Ash1 Trithorax group protein function in Drosophila. Nat Commun 8: 1649.

Kahn, T. G., E. Dorafshan, D. Schultheis, A. Zare, P. Stenberg et al., 2016 Interdependence of PRC1 and PRC2 for recruitment to Polycomb Response Elements. Nucleic Acids Res 44: 10132–10149.

Kharchenko, P. V., A. A. Alekseyenko, Y. B. Schwartz, A. Minoda, N. C. Riddle et al., 2011 Comprehensive analysis of the chromatin landscape in Drosophila melanogaster. Nature 471: 480–485.

Klymenko, T., and J. Muller, 2004 The histone methyltransferases Trithorax and Ash1 prevent transcriptional silencing by Polycomb group proteins. EMBO Rep 5: 373–377.

Kondo, S., and R. Ueda, 2013 Highly improved gene targeting by germline-specific Cas9 expression in Drosophila. Genetics 195: 715–721.

Laprell, F., K. Finkl and J. Muller, 2017 Propagation of Polycomb-repressed chromatin requires sequence-specific recruitment to DNA. Science 356: 85–88.

Lee, Y., E. Yoon, S. Cho, S. Schmahling, J. Muller et al., 2019 Structural Basis of MRG15-Mediated Activation of the ASH1L Histone Methyltransferase by Releasing an Autoinhibitory Loop. Structure 27: 846–852 e843.

Lewis, E. B., 1978 A gene complex controlling segmentation in Drosophila. Nature 276: 565–570.

Li, Z., X. Fu, Y. Wang, R. Liu and Y. He, 2018 Polycomb-mediated gene silencing by the BAH-EMF1 complex in plants. Nat Genet 50: 1254–1261.

McKay, D. J., S. Klusza, T. J. Penke, M. P. Meers, K. P. Curry et al., 2015 Interrogating the function of metazoan histones using engineered gene clusters. Dev Cell 32: 373–386.

Mohan, M., H. M. Herz, E. R. Smith, Y. Zhang, J. Jackson et al., 2011 The COMPASS family of H3K4 methylases in Drosophila. Mol Cell Biol 31: 4310–4318.

Parks, A. L., K. R. Cook, M. Belvin, N. A. Dompe, R. Fawcett et al., 2004 Systematic generation of high-resolution deletion coverage of the Drosophila melanogaster genome. Nat Genet 36: 288–292.

Pengelly, A. R., O. Copur, H. Jackle, A. Herzig and J. Muller, 2013 A histone mutant reproduces the phenotype caused by loss of histone-modifying factor Polycomb. Science 339: 698–699.

Petruk, S., Y. Sedkov, S. Smith, S. Tillib, V. Kraevski et al., 2001 Trithorax and dCBP acting in a complex to maintain expression of a homeotic gene. Science 294: 1331–1334.

Poux, S., B. Horard, C. J. Sigrist and V. Pirrotta, 2002 The Drosophila trithorax protein is a coactivator required to prevent re-establishment of polycomb silencing. Development 129: 2483–2493.

Reha-Krantz, L. J., 1990 Genetic evidence for two protein domains and a potential new activity in bacteriophage T4 DNA polymerase. Genetics 124: 213–220.

Sanchez, R., and M. M. Zhou, 2011 The PHD finger: a versatile epigenome reader. Trends Biochem Sci 36: 364–372.

Savitsky, M., M. Kim, O. Kravchuk and Y. B. Schwartz, 2016 Distinct Roles of Chromatin Insulator Proteins in Control of the Drosophila Bithorax Complex. Genetics 202: 601–617.

Schmahling, S., A. Meiler, Y. Lee, A. Mohammed, K. Finkl et al., 2018 Regulation and function of H3K36 di-methylation by the trithorax-group protein complex AMC. Development 145.

Schmitges, F. W., A. B. Prusty, M. Faty, A. Stutzer, G. M. Lingaraju et al., 2011 Histone methylation by PRC2 is inhibited by active chromatin marks. Mol Cell 42: 330–341.

Schuettengruber, B., H. M. Bourbon, L. Di Croce and G. Cavalli, 2017 Genome Regulation by Polycomb and Trithorax: 70 Years and Counting. Cell 171: 34–57.

Schuettengruber, B., M. Ganapathi, B. Leblanc, M. Portoso, R. Jaschek et al., 2009 Functional anatomy of polycomb and trithorax chromatin landscapes in Drosophila embryos. PLoS Biol 7: e13.

Schwartz, Y. B., T. G. Kahn, P. Stenberg, K. Ohno, R. Bourgon et al., 2010 Alternative epigenetic chromatin states of polycomb target genes. PLoS Genet 6: e1000805.

Schwartz, Y. B., and V. Pirrotta, 2007 Polycomb silencing mechanisms and the management of genomic programmes. Nat Rev Genet 8: 9–22.

Shearn, A., 1989 The ash-1, ash-2 and trithorax genes of Drosophila melanogaster are functionally related. Genetics 121: 517–525.

Smith, S. T., S. Petruk, Y. Sedkov, E. Cho, S. Tillib et al., 2004 Modulation of heat shock gene expression by the TAC1 chromatin-modifying complex. Nat Cell Biol 6: 162–167.

Stassen, M. J., D. Bailey, S. Nelson, V. Chinwalla and P. J. Harte, 1995 The Drosophila trithorax proteins contain a novel variant of the nuclear receptor type DNA binding domain and an ancient conserved motif found in other chromosomal proteins. Mech Dev 52: 209–223.

Steffen, P. A., J. P. Fonseca, C. Ganger, E. Dworschak, T. Kockmann et al., 2013 Quantitative in vivo analysis of chromatin binding of Polycomb and Trithorax group proteins reveals retention of ASH1 on mitotic chromatin. Nucleic Acids Res 41: 5235–5250.

Steneberg, P., and C. Samakovlis, 2001 A novel stop codon readthrough mechanism produces functional Headcase protein in Drosophila trachea. EMBO Rep 2: 593–597.

Tanaka, Y., Z. Katagiri, K. Kawahashi, D. Kioussis and S. Kitajima, 2007 Trithorax-group protein ASH1 methylates histone H3 lysine 36. Gene 397: 161–168.

Tie, F., R. Banerjee, A. R. Saiakhova, B. Howard, K. E. Monteith et al., 2014 Trithorax monomethylates histone H3K4 and interacts directly with CBP to promote H3K27 acetylation and antagonize Polycomb silencing. Development 141: 1129–1139.

Tripoulas, N., D. LaJeunesse, J. Gildea and A. Shearn, 1996 The Drosophila ash1 gene product, which is localized at specific sites on polytene chromosomes, contains a SET domain and a PHD finger. Genetics 143: 913–928.

Tripoulas, N. A., E. Hersperger, D. La Jeunesse and A. Shearn, 1994 Molecular genetic analysis of the Drosophila melanogaster gene absent, small or homeotic discs1 (ash1). Genetics 137: 1027–1038.

Voigt, P., G. LeRoy, W. J. Drury, 3rd, B. M. Zee, J. Son et al., 2012 Asymmetrically modified nucleosomes. Cell 151: 181–193.

Weaver, T. M., E. A. Morrison and C. A. Musselman, 2018 Reading More than Histones: The Prevalence of Nucleic Acid Binding among Reader Domains. Molecules 23.

Yang, N., and R. M. Xu, 2013 Structure and function of the BAH domain in chromatin biology. Crit Rev Biochem Mol Biol 48: 211–221.

Yuan, W., M. Xu, C. Huang, N. Liu, S. Chen et al., 2011 H3K36 methylation antagonizes PRC2-mediated H3K27 methylation. J Biol Chem 286: 7983–7989.

